# ^13^C-SpaceM: Spatial single-cell isotope tracing reveals heterogeneity of *de novo* fatty acid synthesis in cancer

**DOI:** 10.1101/2023.08.18.553810

**Authors:** Elena Buglakova, Måns Ekelöf, Michaela Schwaiger-Haber, Lisa Schlicker, Martijn R. Molenaar, Shahraz Mohammed, Lachlan Stuart, Andreas Eisenbarth, Volker Hilsenstein, Gary J. Patti, Almut Schulze, Marteinn T. Snaebjornsson, Theodore Alexandrov

**Affiliations:** Structural and Computational Biology Unit, European Molecular Biology Laboratory (EMBL), Heidelberg, Germany; Division of Tumor Metabolism and Microenvironment, German Cancer Research Center (DKFZ) and DKFZ-ZMBH Alliance, Heidelberg, Germany; Department of Chemistry, Washington University in St. Louis, St. Louis, MO, USA; Center for Metabolomics and Isotope Tracing, Washington University in St. Louis, St. Louis, MO, USA; Department of Medicine, Washington University in St. Louis, St. Louis, MO, USA; Metabolomics Core Facility, EMBL, Heidelberg, Germany; Molecular Medicine Partnership Unit, EMBL and Heidelberg University, Heidelberg, Germany; BioStudio, BioInnovation Institute, Copenhagen, Denmark

**Keywords:** spatial single-cell metabolomics, stable isotope tracing, single-cell analysis, fatty acid synthesis, acetyl-CoA production, ACLY, hypoxia, IDH-mutant glioma, cancer

## Abstract

Metabolism has emerged as a key factor in homeostasis and disease including cancer. Yet, little is known about the heterogeneity of metabolic activity of cancer cells due to the lack of tools to directly probe it. Here, we present a novel method, ^13^C-SpaceM for spatial single-cell isotope tracing of glucose-dependent *de novo* lipogenesis. The method combines imaging mass spectrometry for spatially-resolved detection of ^13^C_6_-glucose-derived ^13^C label incorporated into esterified fatty acids with microscopy and computational methods for data integration and analysis. We validated ^13^C-SpaceM on a spatially-heterogeneous normoxia-hypoxia model of liver cancer cells. Investigating cultured cells, we revealed single-cell heterogeneity of lipogenic acetyl-CoA pool labelling degree upon ACLY knockdown that is hidden in the bulk analysis and its effect on synthesis of individual fatty acids. Next, we adapted ^13^C-SpaceM to analyze tissue sections of mice harboring isocitrate dehydrogenase (IDH)-mutant gliomas. We found a strong induction of *de novo* fatty acid synthesis in the tumor tissue compared to the surrounding brain. Comparison of fatty acid isotopologue patterns revealed elevated uptake of mono-unsaturated and essential fatty acids in the tumor. Furthermore, our analysis uncovered substantial spatial heterogeneity in the labelling of the lipogenic acetyl-CoA pool indicative of metabolic reprogramming during microenvironmental adaptation. Overall, ^13^C-SpaceM enables novel ways for spatial probing of metabolic activity at the single cell level. Additionally, this methodology provides unprecedented insight into fatty acid uptake, synthesis and modification in normal and cancerous tissues.

## Introduction

Lipids are a highly complex class of hydrophobic biomolecules that are involved in multiple cellular functions. For example, glycerophospholipids and sphingolipids are major structural components of cellular membranes ^1^ while triglycerides stored in lipid droplets provide cells with an additional energy source ^2^. Lipids can also contribute to cellular signaling pathways by functioning as second messengers or lipid mediators ^3-5^. Altered lipid metabolism is a hallmark of many cancers, as it is a prerequisite for rapid growth and proliferation, and promotes survival under nutrient limiting conditions ^6-8^. Altered lipid metabolism in cancer also contributes to oxidative stress resistance and activates oncogenic signaling pathways through the production of lipid mediators ^9^. Lipid metabolism is also affected by different stresses imposed by the tumor microenvironment, such as limited nutrient and oxygen supply ^10^. Therefore, cancer cells need to adapt to these stresses by adopting different routes for the provision of lipids, thus inducing selective metabolic dependencies that could be targeted for cancer therapy.

Most lipids are synthesized from fatty acids, a group of small linear molecules consisting of a carboxylic acid head and an aliphatic tail (Fig. 1a). Fatty acids differ in their length and degree of saturation, i.e. the number of double bonds in the hydrocarbon chain ^11^. Cells in the adult mammalian organism generally take up fatty acids from the blood stream. However, cells of adipose and hepatic tissues as well as various cancers can also generate fatty acids through *de novo* synthesis using acetyl-coenzyme A (acetyl-CoA) as a substrate ^7^. The first step in fatty acid biosynthesis consists of the conversion of acetyl-CoA into malonyl-CoA by acetyl-CoA synthetases (ACC1/2). Malonyl-CoA is then used by fatty acid synthase (FASN), a large enzyme with multiple catalytic centers, for the repeated condensation reactions of two-carbon units, derived from acetyl-CoA, to form palmitate (16:0), a 16-carbon saturated fatty acid. Palmitate can then be further elongated and/or desaturated thus generating a variety of saturated and mono-unsaturated fatty acids that can subsequently be incorporated into the lipidome. The fatty acid composition (or saturation state) of the lipidome is a critical factor for cell survival under metabolically challenging conditions. An example of this is the mono-unsaturated fatty acid oleate which protects cancer cells from lipotoxicity and ER stress ^12-15^, inhibits apoptosis ^16^ and promotes ferroptosis resistance ^17^. Indeed, it has been shown that hypoxic and Ras-transformed cells preferentially take up mono-unsaturated fatty acid to support proliferation and survival ^18^.

**Figure 1.**
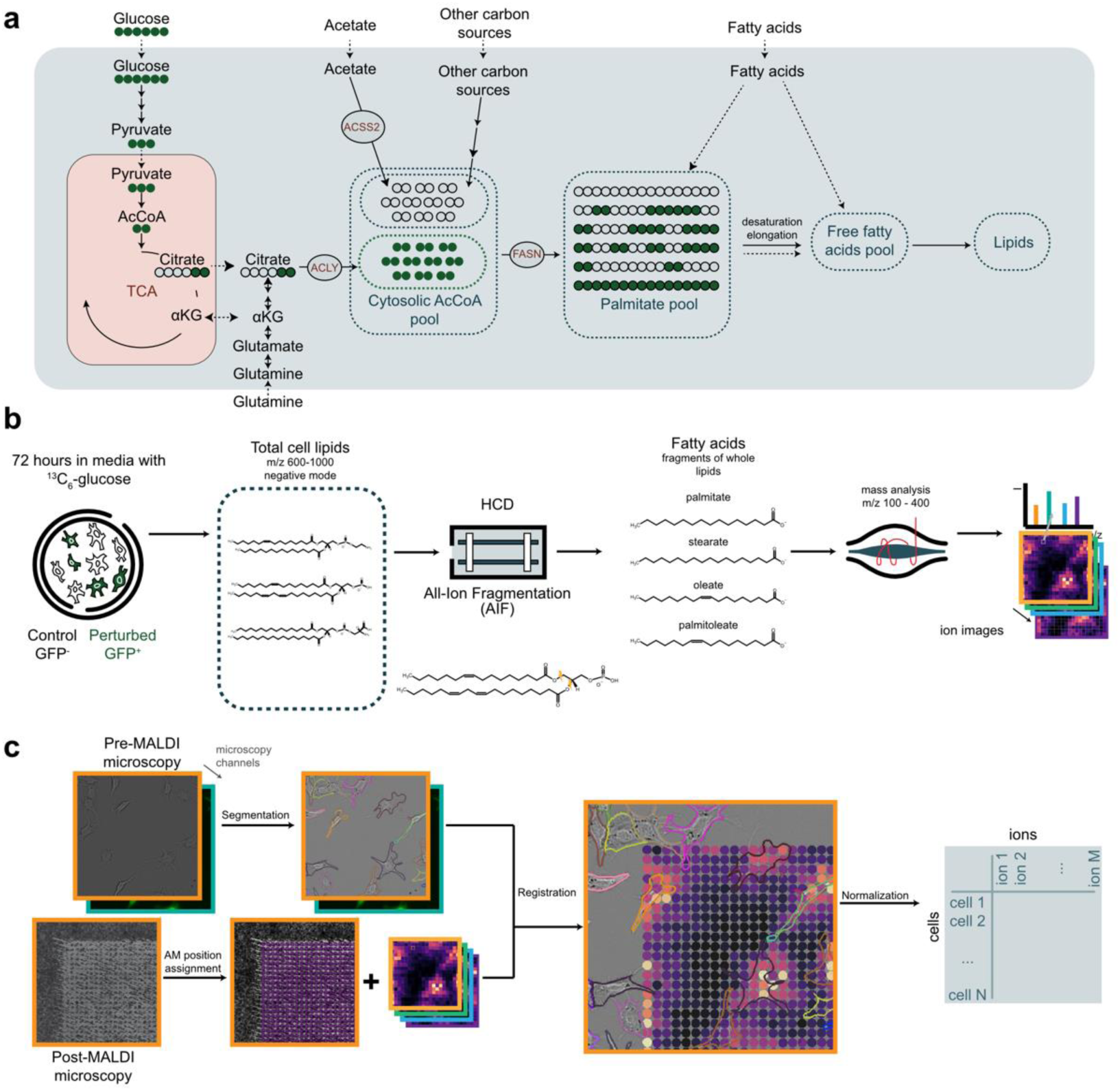
^13^C-SpaceM workflow as applied to interrogate *de novo* fatty acid synthesis. **a)** Generation of cytoplasmic acetyl-CoA and subsequent fatty acid and lipid synthesis from stable-isotope labelled ^13^C_6_-glucose. Glucose-derived pyruvate is metabolized in the mitochondria and exported to the cytoplasm in the form of citrate. ATP-citrate lyase (ACLY) converts citrate to acetyl-CoA, thereby contributing to the cytoplasmic (lipogenic) acetyl-CoA pool. *De novo* fatty acid synthesis by fatty acid synthase (FASN) results in the formation of palmitate (C16:0), which is either further modified or directly incorporated into cellular lipids. Exogenously taken up fatty acids are also incorporated into cellular lipids but are not labelled. **b)** Using all-ion fragmentation imaging mass spectrometry for ^13^C-SpaceM. Cells are grown in medium supplemented with uniformly labelled ^13^C_6_-glucose for 72 hours. The combination of wide-range isolation of parent lipid ions followed by HCD fragmentation and selective isolation of fatty acid fragments allows high-sensitivity measurements of fatty acids incorporated into lipids, conceptually similar to bulk extractions followed by saponification but with the benefit of retained spatial information and the exclusion of free fatty acids. **c)** Integration of microscopy and imaging mass spectrometry in ^13^C-SpaceM to obtain single-cell profiles. Pre-MALDI microscopy and post-MALDI microscopy images containing information about the cell outlines and areas ablated by MALDI-imaging are registered. Ion intensities from MALDI-imaging are assigned to single cells through a normalization procedure.

The cytoplasmic (lipogenic) acetyl-CoA pool used as a substrate for fatty acid synthesis can be derived from different precursors through specific metabolic routes. Glucose-derived carbons provide fuel for the production of citrate by the TCA cycle in the mitochondrial matrix. This citrate is then exported from the mitochondria and used in the cytoplasm by ATP citrate lyase (ACLY) to generate acetyl-CoA. However, the use of glucose as a precursor for fatty acid synthesis is limited under certain conditions, such as hypoxia, in which glucose-derived pyruvate is diverted away from mitochondrial metabolism and converted to lactate. Under these conditions, glutamine can substitute as a precursor for fatty acid synthesis through the reductive carboxylation of glutamate to citrate ^19, 20^. Furthermore, hypoxic cancer cells also use acetate as a precursor for fatty acid biosynthesis through acetyl-CoA synthetase 2 (ACSS2) ^21, 22^.

Besides its function as a substrate for fatty acid synthesis, acetyl-CoA is also needed for the post-translational modification of proteins, most importantly histones for chromatin regulation ^23^. Indeed, both ACLY and ACSS2 have been found to localize to nucleus and modulate histone acetylation and thus affect gene expression ^24, 25^. The modulation of acetyl-CoA metabolism by oncogenic signaling pathways and environmental factors can thereby affect complex gene expression programs in cancer cells. Monitoring the interplay between fatty acid synthesis and acetyl-CoA metabolism therefore provides vital information on the state of cancer cells.

The advent of single-cell tools has revolutionized the analysis of individual cells within heterogenic populations ^26^. The emergence of single-cell transcriptomics and proteomics has provided substantial insights into the single-cell heterogeneity of transcriptional and translational programs yet direct probing of metabolism of single cells has thus far been out of reach due to the lack of capable single-cell metabolomics tools ^27^. It is of particular importance to be able to interrogate fatty acid synthesis and acetyl-CoA metabolism in cancer cells at the single-cell level due to the importance of these processes and the heterogeneity demonstrated for some aspects of metabolism. Different experimental methods have been applied to evaluate the heterogeneity of lipid metabolism in cancer cells. For example, a metabolic sensor based on the malonyl-CoA responsive *Bacillus subtilis* transcriptional repressor, FapR, was developed to determine the activity of FASN in live cells ^28^. Another approach used surface-immobilized dendrimers to evaluate fatty acid uptake on a single cell level ^29^. Furthermore, a recent study applied dielectric barrier discharge ionization (DBDI) on single cells to determine alterations in lipid metabolism upon ACLY inhibition in pancreatic cancer cells ^30^. However, these methods fail to provide broad-spectrum detection of metabolic precursors of fatty acid synthesis and do not allow for sufficient molecular resolution to interrogate acetyl-CoA metabolism.

We previously developed a spatially-resolved mass spectrometry pipeline for the quantification of intracellular metabolites at single-cell resolution ^31^. At the same time, stable-isotope tracing has emerged as a powerful approach to determine *de novo* fatty acid biosynthesis and the contribution of different metabolic precursors to the cytoplasmic acetyl-CoA pool ^21, 32^. We now describe ^13^C-SpaceM, a single-cell method that combines stable isotope tracing with the detection of fatty acids present as acyl-chains in selected lipid classes in a spatially resolved manner, using matrix-assisted laser desorption/ionization (MALDI) in combination with all ion fragmentation (AIF). We applied this method to interrogate the effect of hypoxia on *de novo* fatty acid synthesis in a population of murine liver cancer cells. We showcase it by evaluating the effect of genetic depletion of ACLY on the contribution of glucose to the cytoplasmic acetyl-CoA pool on a single-cell level and by showing the single-cell heterogeneity of the produced effect on the cells. We adapted ^13^C-SpaceM to interrogate tissue sections of mice harboring GL261 glioma cells and could demonstrate spatially-defined alterations in fatty acid uptake, synthesis and modification between glioma tumors and the surrounding normal brain. This spatial analysis revealed activation of *de novo* fatty acid synthesis in tumor tissue and uncovered substantial spatial heterogeneity in acetyl-CoA labelling degree. Overall, the ^13^C-SpaceM method opens novel avenues to interrogate single-cell and spatial heterogeneity in fatty acid synthesis and acetyl-CoA metabolism in cancer.

## Results

### 13C-SpaceM profiles *de novo* fatty acid synthesis of single cells and resolves normoxic and hypoxic metabolic states

The key development presented in this paper is ^13^C-SpaceM, a method for single-cell isotope tracing. ^13^C-SpaceM builds upon SpaceM, a method for single-cell metabolomics which integrates mass spectrometry imaging (MSI) for the detection of stable isotope-labeled metabolites and lipids, and microscopy for assigning imaging MS pixels to individual cells and for quantifying fluorescence and morphometric properties of single cells ^31^. In contrast to the conventional SpaceM method using MS1, in ^13^C-SpaceM we included Higher-Energy Collision-induced Dissociation All Ion Fragmentation (HCD AIF), where lipids in the range of 600-1000 m/z in the negative mode were simultaneously fragmented and then the fragments in the range of 100-400 m/z were detected. This allowed us to selectively interrogate fatty acids that are integrated into cellular lipids. While AIF-MS can be performed in either ion polarity to isolate different groups of common lipid fragments, negative ion mode was chosen for this study as it provides direct detection of fatty acid RCOO-ions. To determine the identities of lipids included in this analysis, we performed MSI with the same mass range and ionization mode without AIF and detected a total of 64 lipid species from several lipid classes, including phosphatidic acids (PA), phosphatidylinositols (PI), phosphatidylethanolamines (PE) and phosphatidylserines (PS). The identities of the majority of these lipids were also confirmed by bulk LC-MS/MS (**Supplementary Table 1 and 2**). Furthermore, the relative abundance of the 11 most abundant fatty acids identified using ^13^C-SpaceM was closely matched by results obtained using bulk MS analysis following saponification and direct infusion (**Supplementary Fig. S1**), confirming the representative coverage of lipids by the method. Registration of microscopy images with mass spectrometry images was done as in conventional SpaceM ^31^ and then after normalization and natural isotope abundance correction the signal was assigned to individual cells (see Methods for details).

We applied ^13^C-SpaceM to interrogate lipid biogenesis through *de novo* fatty acid synthesis, a highly dynamic metabolic process that integrates multiple substrates and contributes to the complexity of cellular lipids (**Fig. 1a**). In order to validate ^13^C-SpaceM in this context, we developed a controlled spatially-heterogeneous model containing co-plated cells previously cultured under either normoxic or hypoxic conditions. We chose this model as hypoxia causes global reprogramming of cellular metabolism, including lipid metabolism. Specifically, hypoxic cells undergo a switch in the use of substrates for the production of cytoplasmic acetyl-CoA used by fatty acid synthesis, by promoting the reductive carboxylation of glutamine to form citrate ^19, 20, 33^ or by the direct use of acetate ^22^. We therefore anticipated that cells cultured under hypoxia should show decreased incorporation of glucose-derived stable-isotope-labeled carbon atoms into cellular lipids compared to cells cultured under normoxic conditions. As fatty acid synthesis exclusively utilizes the cytoplasmic acetyl-CoA pool, analysis of the isotopologues distribution of fatty acids incorporated into cellular lipids allows the assessment of changes in the relative contribution of different substrates to the synthesis of cytoplasmic acetyl-CoA ^21^. The workflow employed by ^13^C-Space allows the quantification of all isotopologues detected for fatty acids released from total lipids isolated at a mass range of m/z 600-1000 in the negative mode using all-ion fragmentation (**Fig. 1b**). Registration of pre- and post-MALDI microscopic images to the MALDI-imaging spectra allows the assignment of ion intensities to individual cells (**Fig. 1c**).

We first applied ^13^C-SpaceM to murine liver cancer cells cultured in normoxia (20% O_2_) and hypoxia (0.5% O_2_) in medium exclusively containing U-^13^C-labeled glucose for 72 hours. This time point was chosen to achieve isotopic steady state for palmitate. Pre-MALDI microscopy images overlaying the brightfield and GFP channels for the co-plated normoxic (GFP^neg^) and hypoxic (GFP^pos^) cells demonstrate equal proportions of both cell populations (**Fig. 2a, first panel**). Cell segmentation and quantification of GFP provided single-cell fluorescence intensities (**Fig. 2a, second panel**). Single-cell isotopologue profiles were constructed by ^13^C-SpaceM for individual fatty acids incorporated into cellular lipids, as shown here for palmitate, the key product of *de novo* fatty acid synthesis. After applying an intensity threshold, the fraction of unlabeled palmitate (M+0) was determined in a pixelated manner and calculated for single cells using ^13^C-SpaceM **(Figure 2a, third and fourth panels**). Underlying mass spectra collected at two representative pixels, one corresponding to a cell cultured under normoxia and another corresponding to a cell cultured under hypoxia showed isotopologues corresponding to labeled palmitate (M>0) only in normoxic cells, while the M+0 peak was predominant in hypoxic cells (**Fig. 2b**). Thus, ^13^C-SpaceM was able to accurately detect differences in isotopologue patterns of palmitate in response to environmental perturbation at the single cell level.

**Figure 2.**
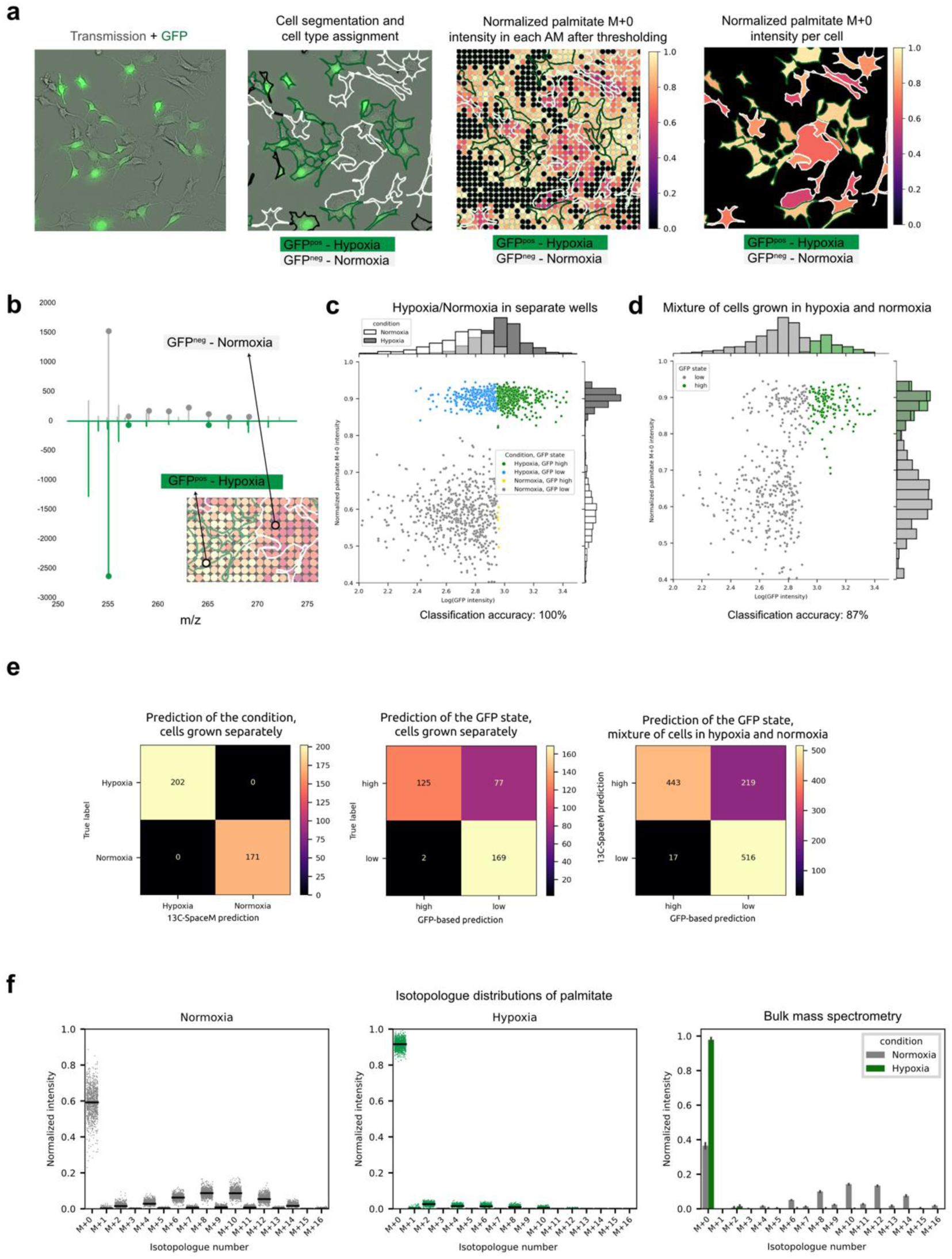
Validating ^13^C-SpaceM by interrogating *de novo* fatty acid synthesis in spatially-heterogeneous normoxia-hypoxia model. **a)** Illustration of microscopy and imaging mass spectrometry data from the model of co-plated primary murine liver cancer cells originally cultured under normoxia (GFP^neg^) and hypoxia (GFP^pos^). The GFP signal was used for discerning the culturing conditions (white and green cell outlines shown normoxic and hypoxic cells, respectively). The normalized intensities of the M+0 isotope of palmitate (representing the fraction of unlabelled palmitate) are shown for MALDI-imaging pixels and as assigned to the single cells. **b)** Mass spectra for individual pixels mapped to cells from a normoxic cell (white outline) and a hypoxic cell (green outline). The peaks corresponding to palmitate isotopologues are marked by grey or green dots. **c)** Discerning cells cultured under normoxia vs. hypoxia using ^13^C-SpaceM for the cells mono-plated for each culturing condition. Scatterplot and histograms show the values of the GFP-reporter (ground truth for telling the condition) and the normalized intensity of the M+0 isotope of palmitate representing the fraction of unlabeled palmitate for single cells. Different colors are used to show cells with different true state label, known from the growth conditions for the particular well, and different cell state assigned using GFP signal. Prediction accuracy refers to the prediction of the true state based on the isotopologue distributions. **d)** Same analysis as in (c) for spatially-heterogeneous co-plated cells from both conditions. In this case accuracy refers to prediction of the cell state assigned using GFP signal. **e)** Confusion matrices for the prediction of culture condition based on ^13^C-SpaceM or GFP in cells grown separately and comparison of the two predictions for co-plated cells. **f)** Comparison of single-cell vs. bulk intensities for the M+0 isotope of palmitate. Single-cell intensities are from the spatially-heterogeneous co-plated model. Bulk intensities are from cells of each mono-cultured condition subjected to total fatty acid analysis by saponification followed by LC-MS. For single-cell data, black lines show average values. For bulk data, data are displayed as mean ± standard deviation across 3 replicates.

In addition to applying ^13^C-SpaceM to the spatially-heterogeneous mix of normoxic and hypoxic cells, we interrogated each condition separately by using mono-plated cultures of the same cells. This was performed to validate that the GFP readout is informative of the condition (GFP^pos^ for hypoxia vs GFP^neg^ for normoxia) and to evaluate whether the isotopologue distribution at the single-cell level can reveal reduced glucose contribution to *de novo* fatty acid synthesis and lipid biogenesis under hypoxia. **Figure 2c** shows a single-cell scatterplot displaying log (GFP intensity) as ground-truth readout characteristic of normoxia (x axis) vs normalized palmitate M+0 intensity calculated using single-cell isotopologue profiles (y axis). Palmitate was chosen for this analysis as it is the immediate product of *de novo* fatty acid synthesis, thus representing the most direct readout of the pathway ^7^. This analysis showed that cells cultured in hypoxia (green) display normalized M+0 intensities close to 0.9, indicating that only a small amount of palmitate was synthesized from glucose. This reflects either an overall low rate of *de novo* fatty acid synthesis or a switch from glucose to other substrates, such as glutamine or acetate ^21^. This is in contrast to cells cultured under normoxia (grey), where the proportion of unlabeled palmitate was substantially lower. Interestingly, cells cultured under normoxia demonstrated higher heterogeneity compared to cells cultured under hypoxia. This heterogeneity could not be attributed to differences in cell size distribution between the two populations (**Supplementary Fig. S1c**) or other morphometric parameters (data not shown). Despite this higher heterogeneity, cells cultured under normoxia displayed ratios of unlabeled palmitate (< 0.8) that were clearly distinct from those of hypoxic cells (> 0.85). This demonstrates that *de novo* fatty acid synthesis is active in all normoxic cells, with a detectable contribution from glucose to the cytoplasmic lipogenic acetyl-CoA pool. We next examined glucose-dependent palmitate labeling at the single-cell level of cells cultured under normoxia and hypoxia using the spatially-heterogeneous co-plating model and GFP as the ground-truth for assigning each cell to either normoxia (GFP^neg^) or hypoxia (GFP^pos^) conditions. Cells with log10 GFP intensity of larger than 3 were considered to be from the hypoxia condition. These cells showed a large fraction of unlabeled palmitate (normalized M+0 intensity around 0.9, **Fig. 2d**), indicating low rates of *de novo* fatty acid synthesis from glucose. However, cells identified as low GFP (log10 GFP intensity less than 3) displayed a mixed phenotype with some cells showing a large fraction of unlabeled palmitate (normalized M+0 intensity >0.8) while others showed evidence for palmitate labelling from glucose (normalized M+0 intensity between 0.4-0.8) (**Fig. 2d**). This can also be seen in the mono-plated scenario (**Fig. 2c**) and is most likely caused by a proportion of the hypoxic cells having lost GFP expression, potentially due to silencing of the promoter in the expression construct.

Overall, we observed a strong similarity in the values measured for the unlabeled palmitate pool in the mono-plated and co-plated scenarios (**Fig. 2c** cf. **Fig. 2d**). Despite this similarity, the separation of the two cell populations based on unlabeled palmitate was less obvious in the co-plated scenario, as a small proportion of cells show intermediate levels of palmitate labelling (normalized M+0 intensities between 0.75-0.88). This potentially indicates an error of estimating the unlabeled palmitate fraction for the hypoxic cells due to co-sampling of neighboring normoxic cells, and the nature of this error (slight increase of intensities of isotopic peaks at M+1 to M+16, thus reducing the fraction of unlabeled palmitate). However, the presence of two distinct clusters (i.e. cells with high levels of M+0 palmitate vs cells with low levels of M+0 palmitate) and the high similarity between the separate and co-plated cells indicates that the effect of this error is small and still allows a clear separation between the two phenotypes.

We therefore quantified the capacity of ^13^C-SpaceM and specifically of the fraction of unlabeled palmitate to discern the two considered phenotypes (normoxia vs. hypoxia) in an unbiased manner. For this, we assigned the conditions to single cells by using the levels of the GFP marker at the threshold of log10 of 3 (with values above this threshold indicating the hypoxic condition). After assigning the conditions in this manner, we used the levels of unlabeled palmitate to classify the cells into the two phenotypes, achieving a classification accuracy of 87%. We furthermore investigated the potential of ^13^C-SpaceM in predicting predicting normoxia vs. hypoxia growth condition based on isotopologue profiles from other fatty acids and not only palmitate (16:0, 16:1, 18:0, 18:1 and 14:0). Using a logistic regression classifier, we could predict the growth condition with perfect accuracy (**Fig. 2e**, left panel). Interestingly, using the GFP fluorescence as the reporter of the growth conditions led to close to 25% false negatives for the same cells. For mixtures of normoxia and hypoxia cells plated together, the GFP prediction was compared to the ^13^C-SpaceM prediction (right panel), with a similar number of false negatives from the GFP (22% of cells). Overall, the high accuracy when using ^13^C-SpaceM validates the capacity of this method to identify the reduced contribution from glucose to palmitate through *de novo* fatty acid synthesis and thus determine the metabolic activity in a spatially-heterogeneous scenario at single-cell resolution. Moreover, the errors of predicting the normoxia vs hypoxia-induced states are likely due to the imperfection of the GFP reporter.

As an additional step of validation, we compared the single-cell isotopologue profiles as provided by ^13^C-SpaceM with bulk measurements of lipids extracted from pooled cells using chloroform/methanol extraction, which allows the isolation of all major phospholipids as well as neutral lipids and ceramides ^34,35^. Lipid extracts were subjected to alkaline hydrolysis (saponification) to release fatty acids and palmitate isotopologue distribution was determined by LC-MS. This comparison showed an overall level of similarity with the pseudo-bulk data for palmitate detected by ^13^C-SpaceM (**Fig. 2f**). However, there was a noticeable difference in the M+0 fraction detected in normoxic cells between bulk and pseudo-bulk analysis (average normalized intensity of 0.6 obtained by ^13^C-SpaceM compared to 0.4 detected in the bulk analysis). One reason for this difference could be that the bulk data represent palmitate released through chemical hydrolysis from a wider variety of lipids. We therefore applied AIF also to bulk LC-MS analysis, using the same mass range (600-1000 m/z) as used in MSI and only using negative mode (**Supplementary Fig. S2a**). This revealed a high similarity in palmitate isotopologue distribution between AIF and saponification, confirming that palmitate molecules released by AIF are representative of the total lipid pool. We also compared the isotopologue distribution of other fatty acids, namely myristate, palmitoleate, stearate and oleate, obtained by MSI and bulk analysis. This showed an overall concordance of the distribution patterns between the two methods (**Supplementary Figure S2b and c**). Importantly, the M+2 isotopologue for stearate and oleate, generated by the elongation of unlabeled precursor, could be detected using both methods. The slightly lower intensity for the M+0 isotopologue obtained using ^13^C-SpaceM could be due to the differences in the interrogated lipid pool (**see Supplementary Table 1**) or the underrepresentation of low abundance isotopologues due to the lower overall sensitivity of MSI.

Summarizing the validation results obtained for the normoxia-hypoxia model, we conclude that ^13^C-SpaceM can profile the contribution of glucose into *de novo* fatty acid synthesis and provide isotopologue distribution reflecting the labeling of palmitate as the key readout of the activity of this pathway, on the single-cell level.

### Quantifying acetyl-CoA pool labeling degree at a single cell level

We next investigated the effect of genetically disrupting components of acetyl-CoA metabolism on fatty acid isotopologue profiles detected using ^13^C-SpaceM. ACLY catalyzes the conversion of cytoplasmic citrate generated from glucose or glutamine into acetyl-CoA (**Fig. 1a**). We therefore engineered cells to express short hairpin RNAs (shRNA) targeting murine ACLY under the control of a doxycycline-inducible promoter. The same promoter also drives expression of a GFP reporter allowing identification of shRNA expressing cells. We used two non-overlapping shRNA sequences (ACLYkd oligo 1 and ACLYkd oligo 2) as well as a non-targeting control. Silencing was achieved by treating cells with 1 µg/ml of doxycycline for 72 hours, with both shRNA sequences resulting in a comparable level of ACLY depletion (**Supplementary Fig. 3a**).

We induced ACLY silencing in cells cultured in the presence of U-^13^C-labeled glucose and interrogated them with ^13^C-SpaceM or bulk MS. **Figure 3a** shows isotopologue distribution for palmitate in the cells expressing non-targeting shRNA and after knockdown of ACLY induced by the two oligos. For the control cells, isotopologue distribution peaks at M+10 in both bulk and single-cell analyses (**Fig. 3a**, left graphs). Pseudo-bulk analysis of ^13^C-SpaceM data showing averages across all single cells indicates a higher M+0 fraction compared to the bulk analysis (normalized peak intensity of 0.6 compared to 0.3). This is in line with the results for the normoxia-hypoxia model in **Figure 2f** and, as discussed earlier, can be explained by differences in the specific lipid pools interrogated by the two methods. ACLY silencing using either oligo induced a marked shift in isotopologue distribution, with lower mass isotopologues being increased (**Fig. 3a**, middle and right graphs). This is in line with our expectation that ACLY silencing reduces the contribution of glucose to the cytoplasmic acetyl-CoA pool.

**Figure 3.**
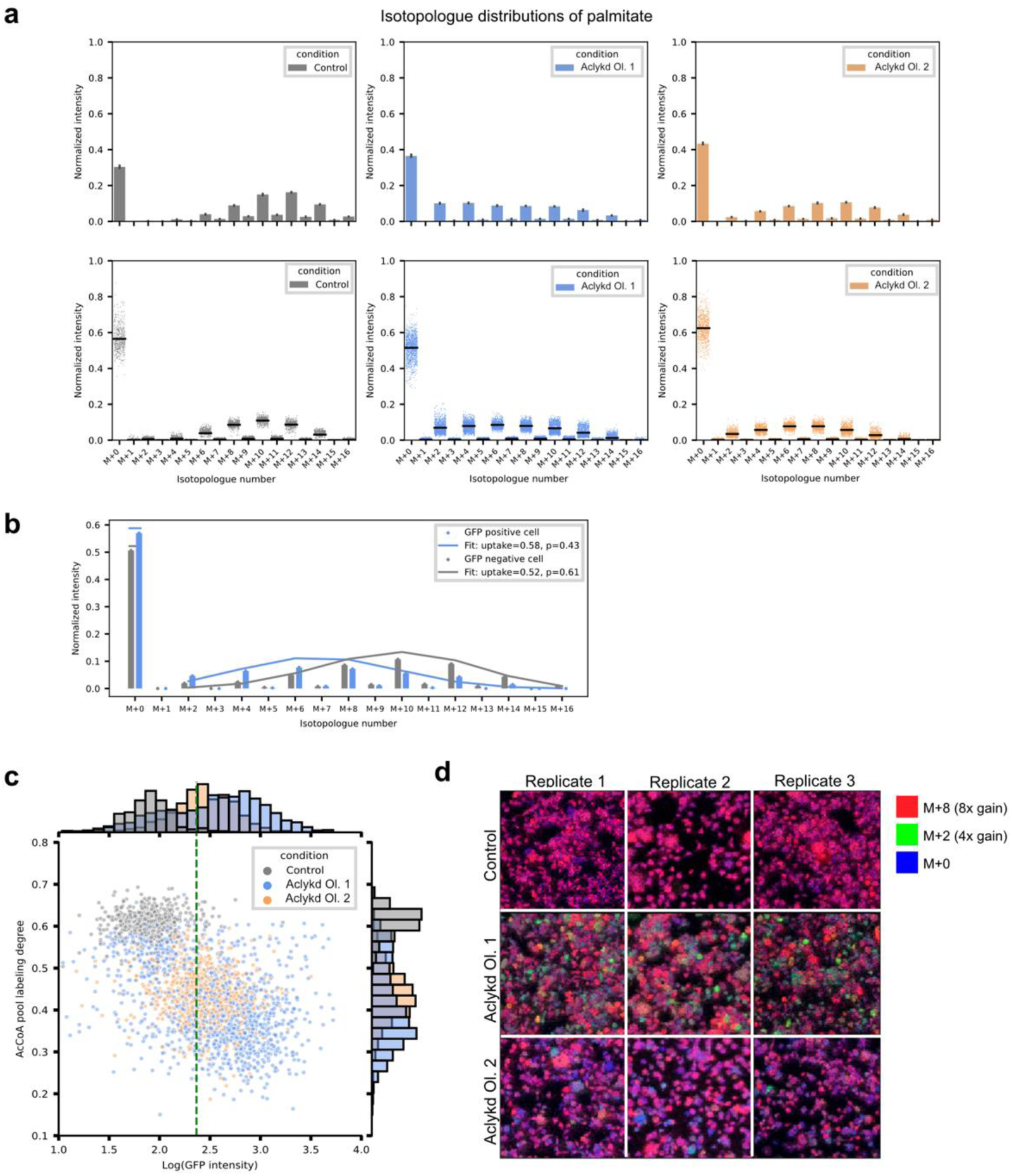
Single-cell quantitative analysis of lipogenic acetyl-CoA production and heterogeneity. **a)** Comparison of bulk and single-cell analysis of isotopologue distribution for palmitate after 72 hours of ACLY gene silencing and culture in the presence of ^13^C_6_-glucose. Bulk data are displayed as mean ± standard deviation across 3 replicates. For single-cell data, black lines show average values. **b)** Normalized single-cell isotopologue distributions for two individual cells, one from the control and the other from ACLYkd oligo 1 (shown as bar plots). Lines show fit of the fatty acid labelling binomial model: horizontal lines for M+0 showing an estimated uptake, and connected lines for M+2i showing (1 - uptake) * binomial(i). Legend shows parameters of the binomial model fit. **c)** Single-cell analysis of the estimated acetyl-CoA pool labelling degree (*p*) as calculated using the fatty acid labelling binominal model for the three conditions: control (grey), ACLYkd oligo 1 (blue), and ACLYkd oligo 2 (blue). Green dashed line shows 95% quantile of the GFP intensity distribution in the control condition, which was used to classify cells as GFP^pos^ vs. GFP^neg^. **d)** Spatial metabolic imaging of *de novo* fatty synthesis for the control, ACLYkd oligo 1, and ACLYkd oligo 2 conditions. Abundance of different palmitate isotopologue peaks are displayed in different channels: M+0 (blue), M+2 (green) and M+8 (red). Each channel is normalized to the total ion count (TIC).

The distribution of the pseudo-bulk intensities across the isotopologues detected after ACLY silencing using oligo 1 suggested the presence of two populations with distinct responses to gene silencing. We therefore used the power of single-cell analysis to deconvolve palmitate labeling in individual cells from the three conditions (control, ACLYkd oligo 1 and ACLYkd oligo 2). We calculated the acetyl-CoA pool labeling degree for each cell by applying a binomial model (described in the methods), with the estimated value *p* quantifying the fraction of labeled acetyl-CoA derived from glucose in the cytosolic acetyl-CoA pool. This model can be applied to each single-cell profile of isotopic intensities of any fatty acid by estimating *p* that leads to best approximation of the data (the number of acetyl-CoA units *n* is equal to half the number of carbon atoms in each fatty acid, e.g. n=8 for palmitate). **Figure 3b** shows examples of such modeling for two cells, a control cell (grey) and an ACLYkd oligo 2 cell (blue). As expected, the ACLYkd cell demonstrates a lower acetyl-CoA pool labeling degree (*p* equal to 0.43 vs. 0.61).

Acetyl-CoA pool labeling degree (*p*) for all single cells in the three populations was plotted against the intensities of the GFP knockdown reporter (**Fig. 3c**). This showed a clear difference between the control cells and cells with ACLY knockdown, with knockdown resulting in lower values of *p* (histogram peak in control at 0.6 compared to 0.35 in ACLYkd oligo 1 and 0.4 in ACLYkd oligo 2). Moreover, the differences between the two ACLY knockdown conditions were also clearly visible. Oligo 1 resulted in a bimodal distribution of the values of *p*, with a visibly higher variance (values ranging from 0.2 to 0.65). A bimodal distribution in single-cell results often indicates the presence of two subpopulations. Notably, one mode of the distribution for the acetyl-CoA pool labeling degree in the ACLYkd oligo 1 condition displayed values for *p* in the range of 0.5-0.6 that overlapped with the values exhibited by control cells (*p* of 0.5-0.7). This can be interpreted as the presence of a subpopulation of "poorly-silenced" cells, where the knockdown was not sufficiently induced. Indeed, ACLYkd oligo 1 cells with high values of *p* (above 0.5) also display GFP intensities similar to those of WT cells, possibly due to the loss of the doxycycline-inducible expression cassette. The second mode of the distribution of the acetyl-CoA pool labeling degree (*p*) for ACLYkd oligo 1 cells had a much lower value (*p* = 0.35). This was clearly below the mode of the distribution of p for ACLYkd oligo 2 cells (*p* = 0.42). This indicates the presence of a “strongly-silenced” subpopulation of ACLYkd oligo 1 cells in which silencing was highly efficient, resulting in a strong reduction of acetyl-CoA labelling. This was contrasted by the results using oligo 2, where the acetyl-CoA pool labelling degree (*p*) as well as GFP reporter intensity displayed more homogenous values. Thus, ^13^C-SpaceM was able to detect heterogeneity in ACLY knockdown cells and identify different sub-populations.

Spatial information provided by ^13^C-SpaceM offers another view at cellular heterogeneity. **Figure 3e** shows ion images for three isotopologue peaks of palmitate (M+0 in blue, M+2 in green, M+8 in red) for the three conditions (control, ACLYkd oligo 1 and ACLYkd oligo 2). The M+2 peak was chosen as the most discriminative for the phenotype of low acetyl-CoA labelling specific to the “strongly-silenced” population among ACLYkd oligo 1 cells. The M+8 peak was chosen as a representative for the control condition where fatty acids contain a high proportion of ^13^C. Thus, the difference between M+2 and M+8 as visible in **Fig. 3a** can serve as an indicator of the relative contribution of glucose to the cytoplasmic acetyl-CoA pool and thus be used to display heterogeneity. The data was acquired with the pixel size of 10 µm with cells having an average area of 550 µm^2^, corresponding to 12 pixels per average cell. Ion images for ACLYkd oligo 1 showed a marked heterogeneity in the intensities of M+2 and M+8 isotopic peaks for palmitate that visualizes the presence of two distinct subpopulations in this condition. Interestingly, the observation that these two subpopulations were spatially-heterogeneous ruled out a batch effect or technical artifacts and indicates a single-cell effect. In contrast, ACLYkd oligo 2 cells showed a more homogenous distribution of palmitate M+8 and an overall lower abundance of the M+2 peak. This degree of single-cell and spatial heterogeneity cannot be revealed through bulk analysis, demonstrating the unique advantages of the ^13^C-SpaceM method.

### 13C-SpaceM differentiates fatty acids by their relative uptake in single-cells

We next considered that at isotopic steady state ^13^C-SpaceM data could be used to determine the relative uptake of different fatty acids as compared to their *de novo* synthesis at single cell level (**Fig. 4c**). For each fatty acid, this was achieved by first performing natural isotope correction, normalizing all isotopologue peaks to the sum of all peaks, approximating isotopologues intensities with the binomial model described in the methods section, and taking the value of the “uptake” from the model. (**Figure 4a, upper plot**) shows single-cell values of the relative uptake for the detected fatty acids: saturated fatty acids (SFAs) myristate, palmitate, stearate as well as mono-unsaturated fatty acids (MUFAs) palmitoleate and oleate, as determined in the control liver cancer cells. Interestingly, the levels of relative uptake at 60%-90% (and thus *de novo* synthesis of 10%-40%) are close to the reported rate of 30% of *de novo* lipogenesis in patients with NAFLD ^36^. The two MUFAs, palmitoleate and oleate, showed a substantially higher relative uptake compared to the corresponding saturated fatty acids palmitate and stearate. This indicates that these cells mostly utilize uptake rather than *de novo* synthesis to obtain mono-unsaturated fatty acids. The results on the average levels were confirmed in the bulk LC-MS-based isotope tracing (**Fig. 4a, lower plot**).

**Figure 4.**
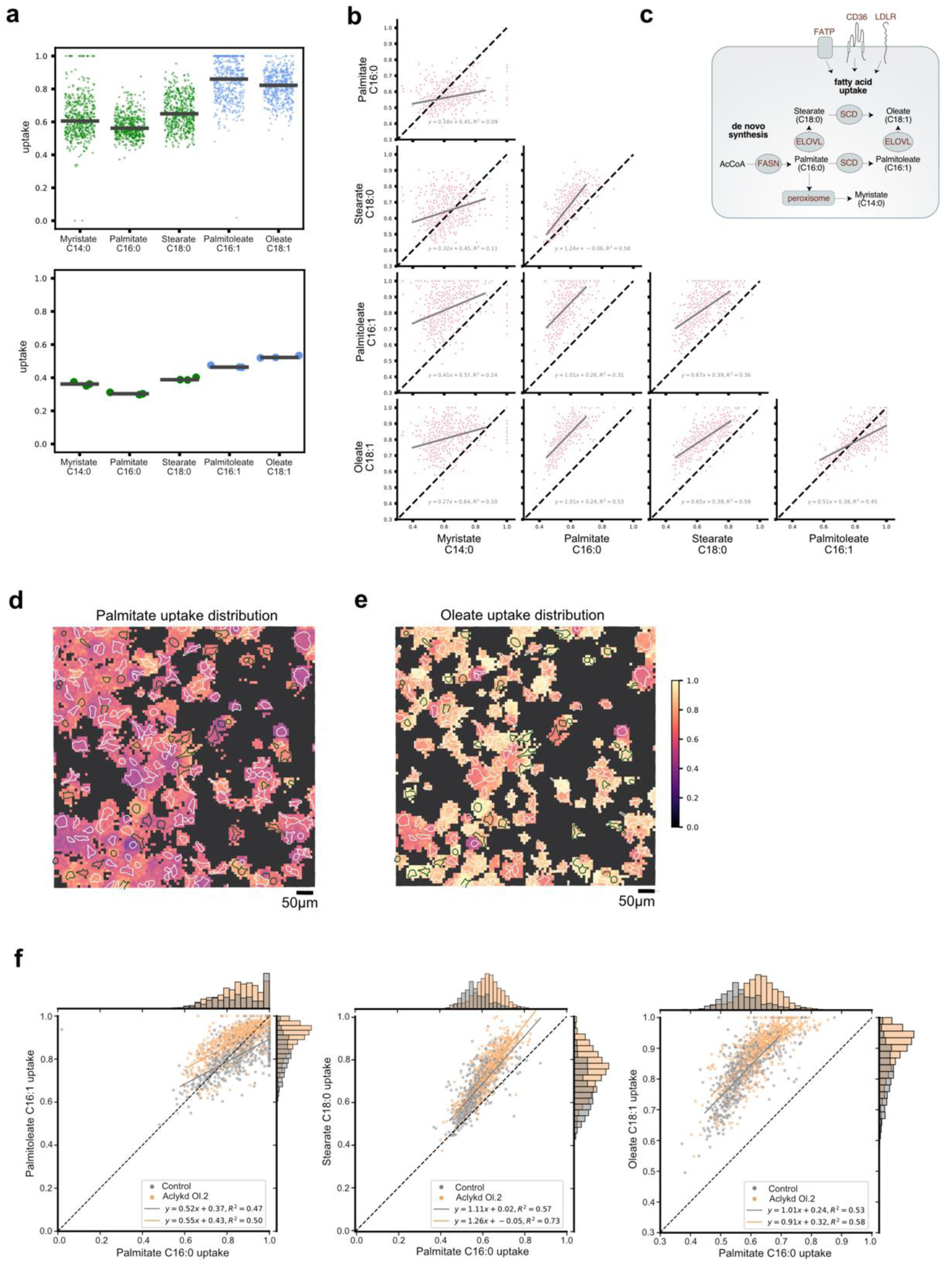
^13^C-SpaceM differentiates fatty acids by their relative uptake in single cells. **a)** Fatty acid uptake in the control cells as determined for myristate, palmitate, palmitoleate, stearate and oleate; saturated fatty acids in green and unsaturated fatty acids in blue. Top: Single-cell data with each point representing a cell, black horizontal lines showing median values. Bottom: Data from the bulk stable isotope tracing. **b)** Correlation of uptake of different fatty acids. Dashed line is the diagonal. For each pair of fatty acids, the linear fit is shown as a grey line, with the fit parameters shown in the legend. **c)** Diagram showing mechanisms involved in fatty acid synthesis, desaturation and uptake. **d)** Spatial metabolic imaging of the palmitate uptake (normalized M+0 fraction) in co-plated cells from control and ACLYkd oligo2 populations. Each square corresponds to one MALDI pixel. Pixels with the total intensity of the fatty acid isotopes below the set limit of detection (1000) are shown in black. Cell outlines are green for GFP^pos^ cells (control) and white for GFP^neg^ cells (Aclykd.oligo2). Pixels are colored according to the normalized M+0 describing the uptake contribution for a given fatty acid. **e)** Same analysis as in (d) for oleate. **f)** Single-cell analysis of changes in fatty acid uptake upon ACLY knockdown for palmitate, palmitoleate, stearate and oleate. Dashed lines are the diagonal. For each condition the linear fit is shown in the corresponding color, with the fit parameters shown in the legend.

We next employed the power of ^13^C-SpaceM to investigate the relationships between the relative uptake of different fatty acids in the same cells. This was done by plotting and analyzing the single-cell values of relative uptake for each fatty acid vs. the others. This analysis showed no correlation of the relative uptake of myristate with any of the other fatty acids (**Fig. 4b, first column**). This lack of correlation is expected, as myristate was reported to be produced by *de novo* FA synthesis only to a small degree. Instead, myristate is mainly generated by the shortening of palmitate via peroxisomal β-oxidation or the elongation of lauric acid ^37^. In contrast, single-cell relative uptake values for palmitate, palmitoleate, stearate and oleate showed strong positive correlations (**Fig. 4b, columns 2-4**). Thus, ^13^C-SpaceM allowed to differentiate fatty acids by their sources specifically by exogenous uptake vs *de novo* synthesis. The lack of correlation for the single-cell uptake values for myristate likely reflects the specific mechanism of production or uptake of this fatty acid.

Next, we investigated the effect of ACLY knockdown on fatty acid uptake. Spatial analysis and visualization demonstrated a strong single-cell heterogeneity in the relative uptake of either palmitate or oleate in a mixed population of control and ACLYkd oligo 2 cells (**Fig. 4d and e**). More detailed and quantified analysis by plotting the single-cell relative uptake values separately for control and ACLYkd oligo 2 showed that ACLY-silenced cells increased the uptake of palmitate (histogram peak at 0.55 in control compared to 0.61 in ACLYkd), with each population displaying a unimodal symmetric distribution (**Fig. 4f**). Oleate uptake, which was already higher compared to palmitate uptake in the control cells, was also increased after ACLY knockdown (histogram peak at 0.85 in control compared to 0.95 in ACLYkd). In addition, we observed an asymmetric (skewed) unimodal distribution of the oleate single-cell relative uptake values for the ACLYkd population, with more cells showing high relative uptake values. This indicates that some cells of the population respond to ACLY silencing by selectively inducing the uptake of mono-unsaturated fatty acids, such as oleate.

### 13C-MSI coupled to AIF reveals intra-tumoral heterogeneity of fatty acid synthesis at the near-single-cell spatial resolution

Our results have shown that combining MALDI imaging MS with AIF to detect esterified fatty acids instead of the lower-abundant free fatty acids substantially increases sensitivity and allows the analysis of isotopologue distribution to assess cytoplasmic acetyl-CoA labelling at the level of single cells. This prompted us to apply this methodology to tumor tissue sections at a near-single-cell spatial resolution to investigate whether metabolic constraints imposed by the tumor microenvironment, such as differential access to nutrients and/or oxygen result in intra-tumoral heterogeneity in terms of glucose contribution to the cytosolic acetyl-CoA pool or the relative contribution of fatty acid uptake vs *de novo* synthesis. We analyzed brain tissue sections from mice that had been orthotopically implanted with GL261 glioma cells expressing mutant isocitrate dehydrogenase (IDH) and red fluorescent protein (RFP). The mice were fed an unlabeled or U-^13^C glucose-containing liquid diet for 48 hours prior to tissue harvest. Sections from the same tissue samples had been analyzed previously using MALDI- and DESI-imaging MS to study the incorporation of glucose-derived carbons into various different metabolites including free (non-esterified) fatty acids ^38^. That analysis revealed spatial differences in isotopologue distribution of free palmitate and stearate between tumor and non-tumor regions of the brain, but did not provide sufficient spatial resolution to fully appreciate intra-tumoral heterogeneity ^38^.

We first applied our methodology to analyze the esterified fatty acid composition in brain sections from tumor-bearing mice that were fed a ^12^C-glucose diet. Comparison of the total ion count (TIC) with brightfield and fluorescent imaging revealed high ion counts throughout the whole brain including the tumor area (**Fig. 5a**). Note that the tumor area (**Fig. 5a**, TIC plot, boxed) was analyzed using a higher resolution (10 µm) as compared to the rest of the tissue section (50 µm). Spatial analysis of different fatty acids revealed a high amount of heterogeneity in fatty acid abundances in the non-tumor bearing hemisphere, with individual structures such as the corpus callosum and anterior commissure identified based on their fatty acid composition alone, with both regions being high in oleate (18:1) and low in palmitate (16:0), stearate (18:0) and arachidonate (20:4) (**Fig. 5b**). Interestingly, while palmitate, oleate, stearate and arachidonate were present at similar levels both in tumors and the surrounding brain, myristate (14:0) and palmitoleate (16:1) were substantially increased in the tumor tissue. Remarkably, the essential fatty acids linoleate (18:2) and alpha/gamma linoleate (18:3) were also selectively higher in the tumor compared to the rest of the brain tissue (**Fig. 5b**).

**Figure 5.**
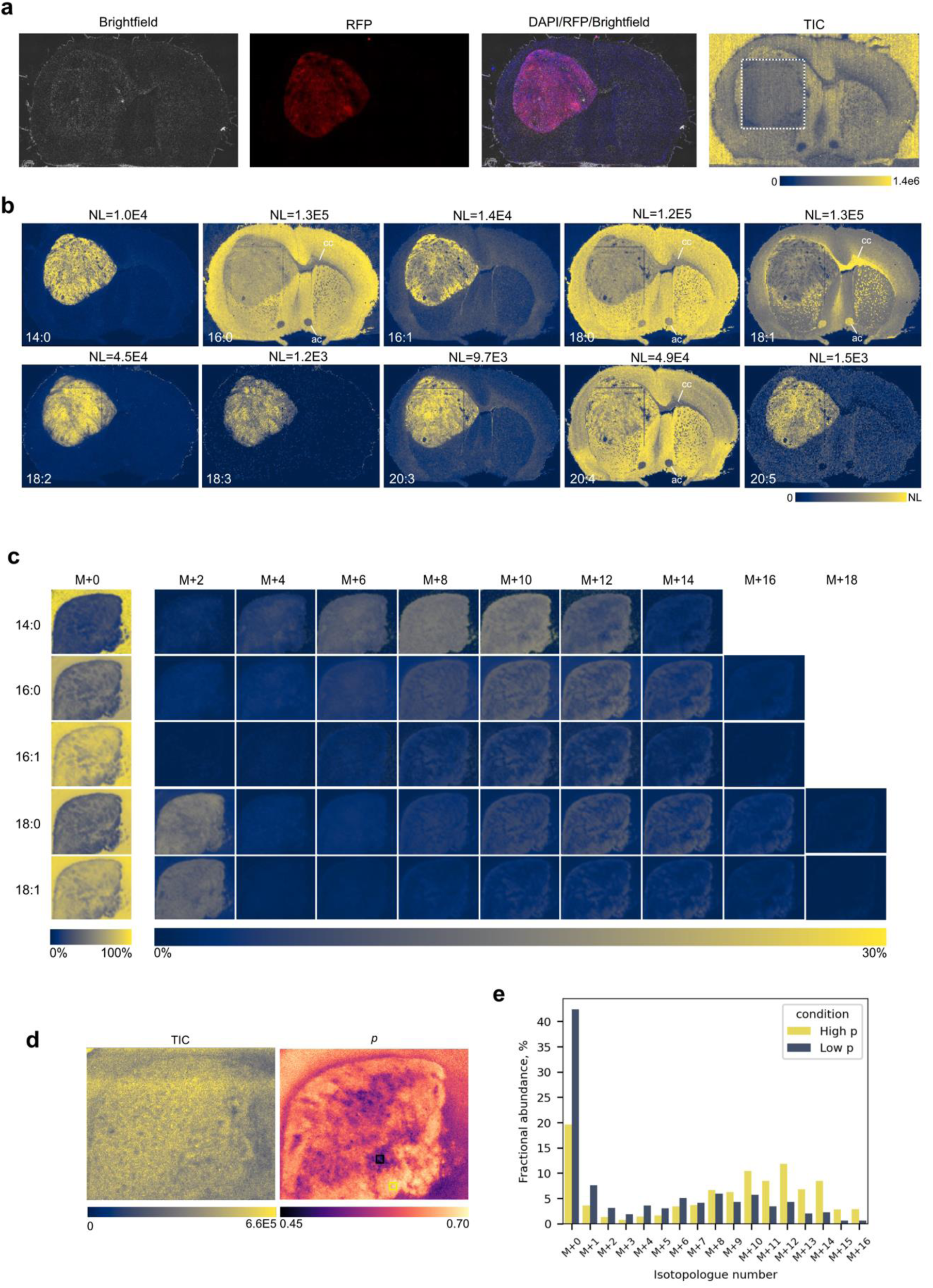
^13^C-SpaceM reveals intra-tumoral heterogeneity of fatty acid synthesis at a near-single-cell resolution. **a)** Panels from left to right: Brightfield image of a cryosection from the brain of a tumor bearing mouse after orthotopic implantation of GL261 glioma cells. Visualization of the RFP expressing glioma cells in the brain. DAPI staining of the cryosection overlapped with the RFP and brightfield channels. Visualization of total ion count (TIC) obtained by MALDI imaging MS and AIF. Square denotes a region where data was acquired at a near-single-cell resolution (10 µm pitch as opposed to 50 µm pitch for the rest of the section). Scale bar ends at the highest point of intensity in the image. **b)** Visualization of the localization of 10 different esterified fatty acids in a section from a tumor bearing brain. NL value (highest signal intensity in the image) for each fatty acid is shown above the image. **c)** Isotopologue fractional images for myristate (14:0), palmitate (16:0), palmitoleate (16:1), stearate (18:0) and oleate (18:1) in tumor tissue (tumor #1) from a mouse orthotopically implanted with GL261 glioma cells and fed with a liquid diet containing U-^13^C_6_-Glucose for 48 hours. Colors represent fraction of each isotopologue of the sum of all isotopologues for each fatty acid. **d)** Visualization of the total ion count (TIC) and the degree of labeling of the cytoplasmic acetyl-CoA pool (*p*) in the tumor. **e)** Isotopologue distributions derived from a high and a low *p* area of the tumor (marked by yellow and black squares in Fig. 5d, respectively).

We next analyzed tissues from tumor bearing mice that had been fed a diet containing U-^13^C glucose for 48 hours and determined the isotopologue distribution of the 5 major esterified fatty acids derived from *de novo* fatty acid synthesis selectively in the tumor tissue (**Fig. 5c and Supplementary Fig. 4a**). The three saturated fatty acids, myristate (14:0), palmitate (16:0) and stearate (18:0) showed a high relative abundance of ^13^C incorporation, with isotopologue distributions peaking at M+10, M+12 and M+14, respectively (**Fig. 5c**). Interestingly, myristate was almost exclusively derived from *de novo* fatty acid synthesis, as the intensity of M+0 was very low compared to the other isotopologues. As myristate is important for the post-translational modification of important signaling proteins ^39^, this finding suggests that glioma tumors may selectively upregulate myristate synthesis to promote their growth. In contrast, the two mono-unsaturated fatty acids, palmitoleate (16:1) and oleate (18:1) showed a higher relative abundance of the M+0 isotopologue (**Fig. 5c and Supplementary Fig. 4b**). In addition, stearate and oleate exhibited a pronounced abundance of the M+2 isotopologue, indicative of elongation from unlabeled precursors (i.e. palmitate and palmitoleate, respectively). This was in line with the previous analysis demonstrating prominent fatty acid elongase activity ^38^.

To analyze this in more detail, we calculated the fraction of all ^13^C isotopologues of the total 1 − [*M* + 0]/ ∑_*n*_[*M* + *n*], representing the fraction of the respective fatty acid that was derived from *de novo* synthesis. This revealed that the majority of palmitate in the tumor was derived from *de novo* fatty acid synthesis (**Supplementary Fig. 4a**). In contrast, the fraction of palmitoleate derived from *de novo* fatty acid synthesis was lower (**Supplementary Fig. 4a**), despite palmitoleate being enriched in the tumor tissue (**Fig. 5b**). This suggests that palmitoleate in the tumor is primarily derived from uptake. Oleate also showed a high labelling degree (**Supplementary Fig. 4a**), but this was mostly caused by the high proportion of the M+2 isotopologue (**Fig. 5c and Supplementary Fig. 4b**) which is formed by elongation of unlabelled palmitoleate. These findings suggest that tumors have limited Δ9-desaturase activity, provided by stearoyl-CoA desaturase (SCD), and rely on the microenvironment for the provision of mono-unsaturated fatty acids (MUFAs).

We next used the isotopologue distribution of palmitate to calculate the fraction of the cytosolic acetyl-CoA pool derived from glucose on a pixel-by-pixel basis, similar to the *in vitro* analysis (**see Fig. 3c**), to generate a spatial representation of acetyl-CoA labelling in the tumor. This revealed a significant degree of spatial intra-tumoral heterogeneity, with glucose contribution to the cytosolic acetyl-CoA pool (*p*) ranging from 0.45 to 0.70 (**Fig. 5d**). It should be noted that the observed variability in *p* was not due to differences in total ion count (TIC), which was quite uniform across the sample (**Fig. 5d**). Close inspection of selected areas with high and low *p* further highlighted the differences in palmitate isotopologue distribution (**Fig. 5e**). Thus, ^13^C-MSI coupled to AIF is capable of determining the labelling degree of the cytosolic acetyl-CoA pool to visualize spatial intra-tumoral metabolic heterogeneity in tissue sections at the near-single-cell resolution.

### 13C-MSI coupled to AIF reveals metabolism of essential fatty acids in glioma

Our finding that the essential fatty acids linoleate (18:2) and alpha/gamma linoleate (18:3) accumulate exclusively in the tumor (**Fig. 5b**) raised the question whether these fatty acids are also processed further, as their derivatives serve as precursors for the synthesis of lipid mediators, such as prostaglandins and leukotrienes that play important roles in cancer ^3^. We therefore analyzed the specific isotopologue distributions of several derivatives of essential fatty acids, namely 20:3, 20:4, 20:5, 22:4 and 22:5 (**Fig. 6a**). We found that the M+2 isotopologue fractions of all except 20:5 were enriched in the tumors. This suggests that glioma tumors not only take up essential fatty acids from the microenvironment but also process these further by elongation and desaturation. Interestingly, the enzymes required for essential fatty acid metabolism, fatty acid elongase 5 (ELOVL5), fatty acid desaturase 1 (FADS1) and fatty acid desaturase 2 (FADS2) are upregulated in low grade glioma (**Fig. 6b and 6c**). Our data also suggest differences in the processing of omega-3 and omega-6 essential fatty acids, as eicosapentaenoic acid (20:5), a PUFA exclusive to the omega-3 branch (**Fig. 6b**), shows no label incorporation (**Fig. 6a**). This suggests that the tumor mostly engages in the processing of omega-6 essential fatty acids, which give rise to pro-inflammatory lipid mediators, such as prostaglandin E2 (PGE_2_).

**Figure 6.**
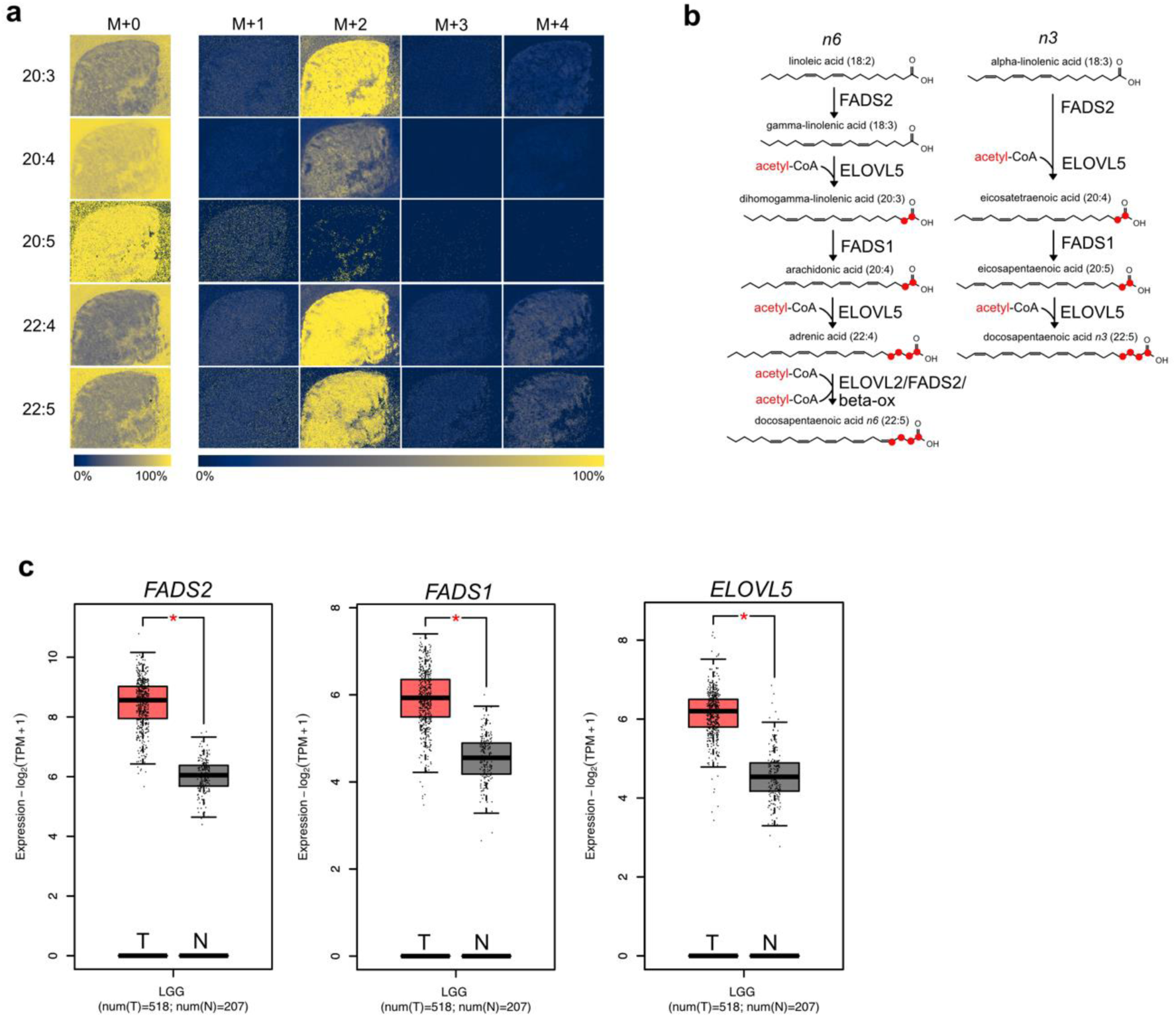
Glioma tumors differentially process *n3* and *n6* essential fatty acids. **a)** Isotopologue distribution for the polyunsaturated fatty acids 20:3, 20:4, 20:5, 22:4 and 22:5. Scale bar shows the fraction of each isotopologue from the sum of the all isotopologues for each fatty acid. **b)** Schematic of the omega-3 and omega-6 polyunsaturated fatty acids biosynthesis pathways. Carbon atoms added from acetyl-CoA by fatty acid elongation are indicated in red. **c)** Comparison of RNA expression for enzymes involved in the processing of essential fatty acids, fatty acid desaturase 1 (FADS1), fatty acid desaturase 2 (FADS2) and fatty acid elongase 5 (ELOVL5) in low grade glioma (LGG) samples from [The Cancer Genome Atlas Program (TCGA)](https://www.cancer.gov/ccg/research/genome-sequencing/tcga) and corresponding normal brain (from The Genotype-Tissue Expression project, GTEx). Analysis was performed using GEPIA2 with a p-value cutoff of 0.01.

Taken together, we have presented ^13^C-SpaceM and ^13^C-MSI coupled to AIF as novel methods for spatial isotope tracing in single cells and in tissue sections at the near-single-cell resolution. We demonstrated how it can be used to interrogate the activity of *de novo* fatty acid synthesis at the single-cell level. Moreover, we expanded its applications to tissue sections where it provided near-single-cell insights into intra-tumoral variations in *de novo* fatty acid synthesis. The key novelty of the method compared to the previously published method SpaceM for spatial single-cell metabolomics is in employing the all-ion-fragmentation MS mode and performing fatty acid isotopologue analysis and modeling. The all-ion-fragmentation is instrumental for the sensitivity required for both single-cell analysis and near-single-cell interrogation of tissue sections. ^13^C-SpaceM can provide detailed insights into the heterogeneity of *de novo* fatty acid synthesis in cancer cells by quantifying heterogeneity of acetyl-CoA metabolism and fatty acid uptake in phenotypically distinct cell populations at the single cell level. The enhanced sensitivity achieved by applying AIF in combination with MSI allows comparable analyses in tissues at the near-single-cell resolution. Concluding, ^13^C-SpaceM addresses the need to investigate the activity of well-characterized metabolic pathways by resolving single-cell and spatial heterogeneity at a high spatial resolution and with a high molecular sensitivity.

## Discussion

The advance of spatial and single-cell omics technologies has revolutionized biology and provided insights into the genetic and phenotypic heterogeneity of cell populations at the levels of tissues, organs and whole organisms ^26^. This is of particular importance in cancer research, as tumors consist of multiple sub-clones of cancer cells as well as a highly diverse population of stromal cells, both of which are exposed to microenvironmental conditions that can change profoundly in a spatial and temporal manner ^40^. Complementing the paradigm of genetic heterogeneity ^41^, there is an emerging acknowledgment of metabolic complexity in cancer ^42^. In addition to genetic and epigenetic alterations, the metabolic state of cancer cells is influenced by non-genetic factors, including the availability of nutrients and oxygen in the microenvironment ^42^. Single-cell transcriptomics helped discover the co-existence of distinct metabolic states in populations of cancer cells and revealed a shift towards oxidative phosphorylation in micro-metastases compared to primary tumors in breast cancer ^43^. Recent technological advances in mass spectrometry, including electrospray ionization (ESI), matrix-assisted laser desorption/ionization (MALDI) or dielectric barrier discharge ionization (DBDI) in combination with advanced computational methods enabled the detection of polar metabolites and lipids from single cells ^30, 31, 44^. While these methods have provided unprecedented insight into the metabolic states of single cells, they fail to capture the dynamic activity of metabolic processes and the degree of flexibility within the metabolic network. At the same time, stable-isotope tracing has emerged as the gold-standard to reveal metabolic activity in cells and tissues in bulk ^45^ and in a spatially-resolved manner ^46^, but was not demonstrated so far in single cells, in particular due to the sensitivity limits. We developed ^13^C-SpaceM, a method to combine stable-isotope tracing with spatial single-cell metabolomics to monitor *de novo* fatty acid biosynthesis in cancer cells at a single-cell resolution. ^13^C-SpaceM builds upon SpaceM ^31^ with major changes including a mass spectrometry method constructed to mimic chemical saponification and subsequent fatty acid detection as well as the novel computational data analysis. In particular, we included a binomial model to determine the labelling degree of the cytoplasmic acetyl-CoA pool from fatty acid isotopologue data for individual cells.

^13^C-SpaceM as employed in this study uses all-ion-fragmentation to release fatty acids from cellular lipids at m/z range of 600-1000 and ionized in the negative mode. This restricts the analysis to fatty acids released from glycerophospholipids, while mostly excluding triglycerides and cardiolipins ^35^. Sphingolipids, such as sphingomyelins, were also most likely excluded due to the CID collision energy applied, which was optimized to achieve maximal sensitivity. Further optimization of collision energy was not possible with the instrument used here, but could be applied in the future to selectively investigate different lipid classes.

We validated the ability of ^13^C-SpaceM to detect changes in glucose-dependent fatty acid synthesis at a single-cell resolution by co-plating cells having two different metabolic states induced by culturing under either normoxia or hypoxia. This model was chosen as hypoxia leads to a global reprogramming of cellular lipid metabolism ^10^. Spatial single-cell isotope tracing by ^13^C-SpaceM clearly separated the two cell populations based on the strong increase in the M+0 fraction of palmitate observed in hypoxic cells. This is expected, as hypoxia reduces the conversion of glucose-derived pyruvate into mitochondrial acetyl-CoA via induction of pyruvate dehydrogenase kinase 1 (PDK1) ^47^, thereby lowering the flux of glucose carbons into the TCA cycle and subsequently into the cytoplasmic (lipogenic) acetyl-CoA pool. At the same time, hypoxia induces a switch in carbon source for lipid synthesis by promoting the reductive carboxylation of glutamine to generate citrate ^19, 20^ and the direct synthesis of acetyl-CoA from acetate ^21, 22^. In addition, hypoxia reduces fatty acid synthesis and stimulates fatty acid uptake ^18, 48^, which also increases the proportion of the palmitate M+0 fraction.

We also assessed the application of ^13^C-SpaceM to monitor the effect of genetic perturbations of the metabolic network in cancer cells. Both bulk and single-cell analysis revealed a marked shift in palmitate isotopologue distribution following silencing of ACLY, the enzyme catalyzing the conversion of citrate to acetyl-CoA. Using a binomial model to quantify fractional labelling from the isotopologue distribution ^21^, we estimated the labelling degree of acetyl-CoA in control and ACLY-silenced single cells. In agreement with previous studies using bulk analysis of palmitate isotopologue distribution in ACLY wildtype vs knockout mouse embryo fibroblasts ^24^, silencing ACLY caused a reduction in the fractional labelling of acetyl-CoA from glucose but only a minor increase in the M+0 fraction of palmitate. This suggests that other substrates, such as acetate, are used to synthesize acetyl-CoA under these conditions ^24^. Our analysis also revealed a substantial degree of heterogeneity in fractional labelling of lipogenic acetyl-CoA within the cell population, particularly in ACLY-silenced cells. This heterogeneity suggested differences in efficiency of the two shRNA sequences used to target ACLY, which were completely obscured in the bulk analysis. As acetyl-CoA is not only used for fatty acid and cholesterol synthesis but is an essential substrate for the post-translational modification of proteins, methods to quantify acetyl-CoA synthesis from different substrates at single-cell level can reveal how genetic or environmental factors drive heterogeneity in cancer.

We also used ^13^C-SpaceM to determine the relative proportion of *de novo* synthesis and uptake of five non-essential fatty acids at single-cell level. While fatty acids can be taken up by cells through passive diffusion, it is generally concluded that uptake functions through receptor-mediated processes ^49^. Interestingly, fatty acid uptake via the scavenging receptor CD36 facilitates metastasis formation by promoting a stem cell phenotype in cancer ^50^. Our single-cell analysis revealed substantial heterogeneity in the relative uptake of different fatty acids within the population of cancer cells. Correlation analysis indicated a higher proportion of uptake of palmitoleate and oleate compared to palmitate, suggesting a higher demand for mono-unsaturated fatty acids. Moreover, ACLY silencing caused a general shift from *de novo* synthesis to fatty acid uptake, which was particular prominent for palmitate and oleate. This is consistent with a recent single-cell lipidomics study observing reduced levels of PC species containing saturated and mono-unsaturated fatty acids upon chemical inhibition of ACLY in pancreatic cancer cells^30^.

Based on the results obtained at a single cell level, we also applied our methodology to brain sections from mice after intracerebral implantation of murine glioma cells. The high sensitivity of the MSI analyses due to using AIF enabled the visualization of different esterified fatty acids in the tumor bearing and non-tumor bearing hemispheres at a near-single-cell resolution. This revealed highly selective partitioning of individual fatty acid species to specific brain structures, which could be further explored for example by integrating our results with spatial transcriptomics data from the brain atlas ^51^ as well as in the future correlative studies combining MSI and other spatial omics.

We also applied our method to mice fed a liquid diet containing U-^13^C glucose for 48 hours to achieve deep labelling of the entire metabolic network, including lipids ^52^. Isotopologue analysis revealed a strong induction of synthesis of saturated fatty acids, myristate, palmitate and stearate, in tumor tissue compared to the surrounding brain, thus confirming previous results obtained in the same model ^38^. This is also in agreement with previous studies indicating that fatty acid synthesis is required for brain metastasis in breast and other cancers ^53, 54^. However, by comparing isotopologue patterns between saturated and mono-unsaturated fatty acids, we found that tumors contained a higher proportion of mono-unsaturated fatty acids derived from uptake. This was highly unexpected, as it has been proposed that mono-unsaturated fatty acids are relatively scarce in in the brain microenvironment, making desaturation an essential metabolic requirement for brain metastasis ^54^. Monitoring metabolism using ^13^C-MSI coupled to AIF thus provides insight into dynamic fatty acid provision in tumor and normal tissues.

In addition, we used the palmitate isotopologue distribution to determine acetyl-CoA labelling in tumor tissue in a spatial manner. This revealed substantial spatial heterogeneity, most likely caused by differences in the local availability of nutrients and oxygen in different tumor areas. The observed differences in palmitate isotopologue pattern, increase in M+0 and shift towards isotopolgues with lower mass, were remarkably similar to those observed after exposure of cancer cells to experimental hypoxia and after silencing of ACLY. Interestingly, it has been shown that ACLY promotes cell migration in glioblastoma through mechanisms in involving histone modification ^55^. Moreover, a recent spatial transcriptomics and proteomics study defined hypoxia as a major driver of long-range tissue organization in glioma ^56^. It will be highly interesting to integrate ^13^C-SpaceM data with other spatial approaches to obtain deeper insight into mechanism of metabolic heterogeneity in cancer.

Finally, we applied our methodology to analyze the metabolism of essential fatty acids. Remarkably, tumors showed strong evidence for the elongation and desaturation of omega-6 essential fatty acids, indicating that these fatty acids could play an important function in glioma biology. This agrees with the elevated expression observed in human low-grade glioma of FADS1, FADS2 and ELOLV5, the enzymes responsible for the metabolism of omega-6 essential fatty acids. Furthermore, we found that tumors obtained a substantial amount of arachidonate from uptake, in addition to their active omega-6 metabolism, most likely as this fatty acid is available in large amounts in the brain. This suggests that tumors have a high demand for arachidonate, potentially for the synthesis of pro-inflammatory lipid mediators, including prostaglandins and thromboxanes, which drive rapid proliferation and immune evasion ^3^. Altered fatty acid metabolism is a hallmark of glioma, including glioblastoma ^57^, and an increase in the ratio of omega-3 and omega-6 fatty acids has been described as a marker of aggressive disease ^58^.

Together, our results present ^13^C-SpaceM as a method for spatial single-cell isotope tracing able to monitor *de novo* fatty acid synthesis, production of lipogenic acetyl-CoA and fatty acid uptake of heterogenous cell populations exposed to genetic and environmental perturbations. Furthermore, ^13^C-SpaceM can be used to obtain deep insight into fatty acid metabolism of normal and cancer tissue. Through additional modifications of this approach, for example by applying different metabolic tracers or by restricting collision-induced dissociation to individual lipid classes, ^13^C-SpaceM can also provide more specific information about lipid metabolism at a single-cell resolution. Furthermore, by detecting other metabolite classes with the underlying MALDI-imaging mass spectrometry as already demonstrated ^46^, ^13^C-SpaceM can be extended for stable isotope tracing of other metabolic pathways thus opening avenues towards revealing and understanding single-cell heterogeneity of metabolic activity. Finally, results obtained with ^13^C-SpaceM can also be integrated with other spatial omics data, such as transcriptomics or proteomics, to obtain deeper insight into the metabolic programs of cells and tissues.

## Material and Methods

### Cell culture for the normoxia-hypoxia model

Primary murine liver cancer cells derived from a Myc- and Akt-driven tumor model (*Myc*^OE^; *Akt*^Myr^; *Tp53*^-/-^) ^15^ were a gift from Daniel Dauch (University of Tübingen). Cells were cultured in glucose-free DMEM (Sigma Aldrich) supplemented with 1 mM acetate, 2 mM glutamine and 10% dialyzed FCS after addition of 25 mM of either ^12^C-glucose (Sigma) or U-^13^C-glucose (Cambridge Isotopes) in a 37°C incubator with 5% CO_2_ either under normoxia (20% O_2_) or hypoxia (0.5% O_2_) for 72 hours. The time point of 72 hours was chosen to make sure that isotopic steady state for palmitate had been reached. Hypoxic conditions (0.5% O_2_) were induced in a hypoxia workstation (H35, Don Whitley). The cells cultured under hypoxic conditions were modified to express Green Fluorescent Protein (GFP) as a marker. For co-plating experiments, the normoxic and hypoxic cells were detached using trypsin, mixed using 10,000 cells of each condition, plated on the same glass slide and allowed to attach for 3 hours before fixation.

### ACLY knockdown

For the ACLY knockdown, we used the same primary murine liver cancer cells as in the hypoxia model. shRNA sequences targeting murine ACLY or non-targeting controls were cloned into LT3-GEPIR (Addgene), the shRNA sequences used were (TGCTGTTGACAGTGAGCGACCGCAGCAAAGATGTTCAGTATAGTGAAGCCACAGATGTATACTGAACATCTTTGCTGCGGCTGCCTACTGCCTCGGA), (TGCTGTTGACAGTGAGCGAACCAGTGTCTACTTATGTCAATAGTGAAGCCACAGATGTATTGACATAAGTAGACACTGGTCTGCCTACTGCCTCGGA), (TGCTGTTGACAGTGAGCGCAGGAATTATAATGCTTATCTATAGTGAAGCCACAGATGTATAGATAAGCATTATAATTCCTATGCCTACTGCCTCGGA) for ACLYkd oligo 1, oligo 2 and nontargeting control, respectively. Lentiviral particles were produced in HEK293 cells after transient transfection of the packaging vectors psPAX.2 and pMD.G2 (Addgene #12260 and #12259). After viral transduction, liver cancer cells were selected with puromycin and used at low passage. Induction of shRNA expression was achieved by treating cells with 1µg/ml doxycycline (Sigma) for 72 hours. For stable isotope tracing, cells were cultured in glucose-free DMEM supplemented with U^13^C_6_-glucose (Cambridge Isotope laboratory, Tewksbury, Massachusetts) and 1 mM acetate for 72 hrs.

### Analysis of fatty acids and lipids using liquid-chromatography coupled with mass spectrometry (LC-MS)

For bulk LC-MS, cells were washed with cold 154 mM ammonium acetate, snap frozen in liquid nitrogen and harvested in methanol/H_2_O (80/20, v/v) with added standards (For fatty acids: 10 µL of 100 µM Palmitate-2,2-D2 (Eurisotop: DLM-1153-0)/ 1 x 10E6 cells For lipids (SPLASH LIPIDOMIX, Avanti Polar Lipids, 330707-1EA) 10 μl/sample). Subsequently, 30 μL 0.2 M HCl, 2 x 100 μL CHCl_3_, 2 x 100 μL H_2_O was then added with vortexing in between. The suspension was centrifuged at 16,000 g for 5 min at room temperature, the lower lipid phase was then washed with synthetic polar phase (CH_3_Cl/Methanol/H_2_O, 58/33/8, v/v/v) and evaporated to dryness under N_2_ at +45°C. For lipidomics the samples were respuspended and subjected to LC-MS analysis. For fatty acid analysis lipid extract was saponified by resuspension in Methanol/H_2_O (80/20, v/v) containing 0.3 M KOH, heating at +80°C for 1 h and washed twice with 0.5 mL hexane. After addition of 50 μL formic acid, fatty acids were subsequently extracted twice with 0.5 mL hexane and evaporated to dryness under N_2_ at +45°C. For LC/MS analysis the fatty acids, were dissolved in 100 μL isopropanol and 5 μL of each sample was applied to a C8 column (Accucore C8 column, 2.6 µm particle size, 50 x 2.1 mm, Thermo Fisher Scientific) at +40°C, with mobile phase A consisting of 0.1% formic acid in CH_3_CN/H_2_O (10/90, v/v), and solvent B consisting of 0.1% formic acid in CH_3_CN/H_2_O (90/10, v/v). The flow rate was maintained at 350 μL/min and eluent was directed to the ESI source of a mass spectrometer from 3 min to 27 min after sample injection. MS analysis was performed on a Q Exactive Plus Orbitrap mass spectrometer (Thermo Fisher Scientific) applying the following settings: Sheath gas, 30; spray voltage, 2.6 kV. Capillary temperature +320°C, aux gas heater temperature: +120°C and S-lens voltage was 55. A full scan range from 150 to 460 (m/z) in negative ion mode was used. The resolution was set at 70,000. The maximum injection time was 100 ms with an AGC-target of 1E6. For lipidomic analysis, lipids were separated on a C8 column (Accucore C8 column, 2.6 µm particle size, 50 x 2.1 mm, Thermo Fisher Scientific) mounted on an Ulitmate 3000 HPLC (Thermo Fisher Scientific) and heated to 40°C. The mobile phase buffer A consisted of 0.1% formic acid in CH3CN/H2O (10/90, v/v) and buffer B consisted of 0.1% formic acid in CH3CN/IPOH/H2O (45/45/10, v/v/v). After injection of 3 µl lipid sample, 20% solvent B were maintained for 2 minutes, followed by a linear increase to 99.5% B within 5 minutes, which was maintained for 27 minutes. After returning to 20% B within 1 minutes, the column was re-equilibrated at 20% B for 5 minutes, resulting in a total run time of 40 minutes. The flow rate was maintained at 350 µl/min and the eluent was directed to the ESI source of the QE Plus from 2 to 35 minutes. MS analysis was performed on a Q Exactive Plus mass spectrometer (Thermo Fisher Scientific) applying the following settings: Scan settings: Scan range – 200-1600 m/z in full MS mode with switching polarities (neg/pos) and data-dependent fragmentation; Resolution – 70,000, AGC target – 1E6; Max. injection time – 50 ms. HESI source parameters: Sheat gas – 30; Aux gas – 10; Sweep gas – 3; Spray voltage – 2.5 kV; Capillary temperature – 320°C; S-lens RF level – 55.0; Aux gas heater temperature – 55°C; Fragmentation settings: Resolution – 17,500; AGC target – 1E5; Max. injection time –50 ms. Peaks corresponding to the calculated fatty acid or lipid masses (±5 ppm) were integrated using El-Maven (https://resources.elucidata.io/elmaven) and correction for natural ^13^C isotopic abundance was done using IsoCorrectoR ^59^.

### Analysis of fatty acid distribution and isotopologue distribution by direct infusion MS

For the analysis of fatty acid distribution and isotopologue distribution using direct infusion of saponified fatty acids harvested as described above were resuspended in 50:50 solution A and B (A: H2O/ACN/Isopropanol 2/10/88, B: H2O/ACN 60/40 both with10 mM ammonium acetate). Samples were injected by direct infusion at 20 μL/min for two minutes whereas a spectra was acquired for a total of 8 minutes (3 minutes before and after injection to establish baseline) using the following scan parameters, Full MS scan range of 69-1000 m/z, AGC target 1e6, resolution of 70,000 in negative mode, maximum injection time was 100 ms. HESI source parameters were as follows, sheath gas flow rate 6, spray voltage 3.20 kV, S-lens RF level 50 and aux gas temperature 120°C. For AIF, samples were not saponified and lipid extract was resuspended the same as above. All parameters were the same except scan type was AIF, precursor masses were collected in the range of 600-1000 m/z, scan range was 200-350 m/z, fragmentation was carried out using CE of 50, maximum injection time was 200 ms.

### Microscopy

After cell culturing, the cells were washed with PBS, fixed using Histofix (Roth, 87.3) for 10 minutes, stained using DAPI, washed 3x in PBS and then desiccated in a Lab Companion Cabinet Vacuum Desiccator for 30 min at room temperature and −0.08 MPa. Pre-MALDI brightfield and fluorescent microscopy (620 and 460 nm) images were obtained with a DS-Qi2 camera (Nikon Instruments) with a Plan Fluor 10x (numerical aperture (NA) of 0.30) objective (Nikon Instruments) mounted on a Ti-E inverted microscope (Nikon Instruments). The pixel size was 0.64 μm. For the hypoxia experiment, additional pre-desiccation microscopy images were collected. Rigid registration of pre-desiccation and post-desiccation microscopy images was performed using the Affinder plugin in Napari ^60^ (https://www.napari-hub.org/plugins/affinder). The cells were imaged in bright-field microscopy after MALDI-imaging, using the same microscopy setup and parameters.

### Microscopy image analysis

Cells in the bright-field channel of pre-MALDI microscopy were segmented using a custom Cellpose model ^61^ trained on manually annotated fragments of images. Since the primary cell culture contains cells of varying size, segmentation for the largest cells had to be manually corrected. Ablation marks were detected in the post-MALDI microscopy by manually fitting a grid of circular shapes. Registration of the pre- and post-MALDI images was done using the SpaceM software tool as previously published ^31^.

### MALDI imaging of single-cell samples

The MALDI matrix 1,5-diaminonaphthalene (DAN) was applied to a surface density of 3 μg/μm^2^ immediately before MALDI analysis using a TM sprayer (HTX Technologies, Chapel Hill, NC, USA). Atmospheric pressure MALDI imaging was performed using an AP-SMALDI5 ion source (TransMIT GmbH, Gießen, Germany) coupled to a Q Exactive Plus Orbitrap mass spectrometer (Thermo Fisher Scientific). Raster pitch was 10x10 μm, with the laser attenuator angle set to 33°. The mass spectrometer method used was set up as AIF in negative ion mode, with an isolation range of 600-1000 m/z and a scan range of 100-400 m/z at 140,000 resolution with 500 ms maximum ion injection time. Fragmentation was performed using higher-energy collisional dissociation (HCD) at a normalized collision energy (NCE) of 25. This energy level was determined by a manual stepped collision energy experiment, where we saw no improvement in fatty acid signal/noise by going as high as 45 NCE.

### SpaceM analysis

The ^13^C-SpaceM method is based on SpaceM described in detail earlier ^31^. The key differences are in using AIF (see “MALDI imaging of single-cell samples”) and processing obtained profiles (see the respective section of Methods). Briefly, SpaceM integrates microscopy images and the MALDI images by detecting the MALDI ablation marks, overlaying them with the segmented cells and performing ablation marks-cells deconvolution by applying a mathematical formula. This results in single-cell profiles of molecules detected by the MALDI imaging mass spectrometry.

#### Constructing single-cell isotopologue profiles

MALDI images for unlabeled samples were annotated with METASPACE ^62^ at 5% FDR to determine the most abundant detected fatty acids. For each fatty acid, intensities corresponding to theoretical isotopologue peak masses were extracted. Raw intensities were normalized for natural isotope abundance using IsoCor ^63^, after which every isotopologue distribution was normalized by its sum. Ablation marks with a total raw intensity of less than 200 for a given fatty acid were removed from the analysis. Ablation marks which had at least 30% overlap with the cell mask were considered intracellular. After normalizing spectra in each pixel, the median of values in the intracellular ablation marks was assigned as a single-cell readout for each isotopologue peak. The resulting single-cell isotopologue distribution for every fatty acid was normalized by its sum again. Scanpy v.1.8.1 ^64^ was used for all single-cell data analysis.

#### Calculation of single-cell features

For each well of a multi-well slide, the median intensity of the GFP channel outside of the cell masks was subtracted from the image, and the logarithm of the maximum GFP intensity inside the cell mask was used to characterize single-cell GFP signal. Both in normoxia/hypoxia and in ACLYkd experiments, the 95^th^ percentile of the GFP signal in the GFP-negative wells was used as a threshold to assign cells to either GFP-positive or GFP-negative state.

In addition to the normalized isotopologue peak intensities, a binomial model for the fatty acid synthesis described in ^32^ was used. Fatty acids are synthesized by randomly taking 2-carbon monomers from the cytosolic acetyl-CoA pool. Therefore, if the fraction of labeled monomers equals *p*, the probability that a given fatty acid has 2i labeled carbons, can be described with the following model:

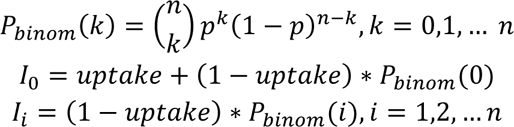

where uptake is the fraction of the unlabeled FA directly taken from the media, *p* is the labeling degree of cytosolic acetyl-CoA pool, and n is the number of acetyl-CoA molecules used for the synthesis (number of carbons in the fatty acid / 2). Longer fatty acids can be synthesized both using labeled palmitate made by the cell *de novo* and unlabeled palmitate, therefore we used a modified model for stearate and oleate (C18):

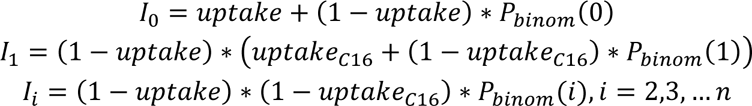

where *uptake_C16_*is the fraction of palmitate (C16) taken from the medium.

Fitting a binomial model to the single-cell isotopologue distributions allows summarizing them as two numbers: uptake, which characterizes the fraction of the fatty acid which was not synthesized *de novo* but directly taken from the medium, and acetyl-CoA pool labeling degree *p*, which describes relative contribution of the labeled substrate to the fatty acid synthesis compared to other carbon substrates consumed by the cell. The limitation of this method is that if overall substrate usage is very low, such as in the hypoxia/normoxia experiment, it becomes impossible to reliably fit a binomial distribution and estimate the uptake fraction, therefore it was only used for the ACLYkd experiment data analysis.

### Orthotopic glioma mouse model

Tissue sections were prepared from mouse brains used for a previous study ^38^. The experiments were approved by the Institutional Animal Care and Use Committee at Washington University (assurance number A338101, protocol 19-0930 and 22-0304) and were performed in accordance with the recommendations in the Guide for the Care and Use of Laboratory Animals of the NIH. In brief, murine glioma cells (GL261-RFP, transduced with IDH1 R132H) were implanted into female mice (C57BL/6J, 8 weeks old). After 8 days, mice were fed a liquid diet containing unlabeled glucose or U-^13^C-labeled glucose (Cambridge Isotope Laboratories) ad libitum for 48 h as previously described ^52 38^. Brains were embedded in 5% wt. carboxymethyl cellulose in water and stored at −80 °C. 10 µm thick sections were collected on Superfrost Plus slides (Thermo Fisher Scientific), dried under vacuum, stored at −80 °C, and shipped on dry-ice. Serial 10 µm thick tissue sections were mounted on Superfrost Plus slides in Fluoroshield mounting medium with DAPI (aqueous, Abcam, USA) and used for fluorescence microscopy as described previously^38^.

### MALDI tissue imaging

Imaging of brain sections was performed using the same sample preparation method and instrumentation described for single cells. For each whole brain section, one image of the tumor and immediate environment was first acquired at 10x10 µm pitch, followed by an image of the rest of the tissue section at 50 µm pitch. The laser attenuation angle was set to 32°, the isolation range was 600-1600 m/z, product scan range 100-600 m/z, and the normalized collision energy set to 30.

### Data analysis for tissue imaging

Each tissue image was converted to the imzML format and annotated with METASPACE ^62^ in the same way as the single-cell data. Compared to processing data from the cells, no cell segmentation was performed and all data analysis was performed on single MALDI pixels (with the pitch of 10 µm within the tumor area and with 50 µm within the rest of the tissue section). The same normalization and AcCoA pool labelling degree modelling was applied as for the cells, and figures were exported as spatial heatmaps using the same custom script as for single-cell data

## Availability of Data and Materials

All MALDI data in this paper is available at https://metaspace2020.eu/project/buglakova-2023. The code for producing the results and figures is available at https://github.com/Buglakova/13C-SpaceM.

## Acknowledgements

We would like to thank I. Germer and AB. Chaves-Filho for technical assistance. We also thank L. Zender and D. Dauch for providing murine liver cancer cell lines (University Hospital Tübingen). A.S and M.T.S would like to acknowledge funding from the German Research Foundation (DFG, FOR-2314 and SCHU 2670-1/2). G.J.P would like to acknowledge the National Institutes of Health grant R35 ES028365. T.A. acknowledges funding from ERC (CoG grant agreement No. 773089, PoC No. 101101077), SNF (PROMETEX), and MJFF.

## Author contributions

TA, EB, MTS and AS conceptualised the study and experimental design. MTS, EB, ME, LS, MRM, SM, LS, AE and VH conducted experiments and analysed data. MSH and GJP designed the in vivo labeling experiments and prepared the in vivo samples. TA, EB, ME, MTS and AS wrote the manuscript with critical input from all authors.

## Supplementary Table legends

**Supplementary Table 1:**

List of masses detected in the murine HCC (*Myc*^OE^; *Akt*^Myr^; *Tp53*^-/-^) cell line using single cell MSI (m/z 600-1000) and automatically annotated using Metaspace.

**Supplementary Table 2:**

List of lipids identified in the murine HCC (*Myc*^OE^; *Akt*^Myr^; *Tp53*^-/-^) cell line using LC-MS/MS.

## Supplementary Figures and Legends

**Figure S1 (accompanying Figure 2):**
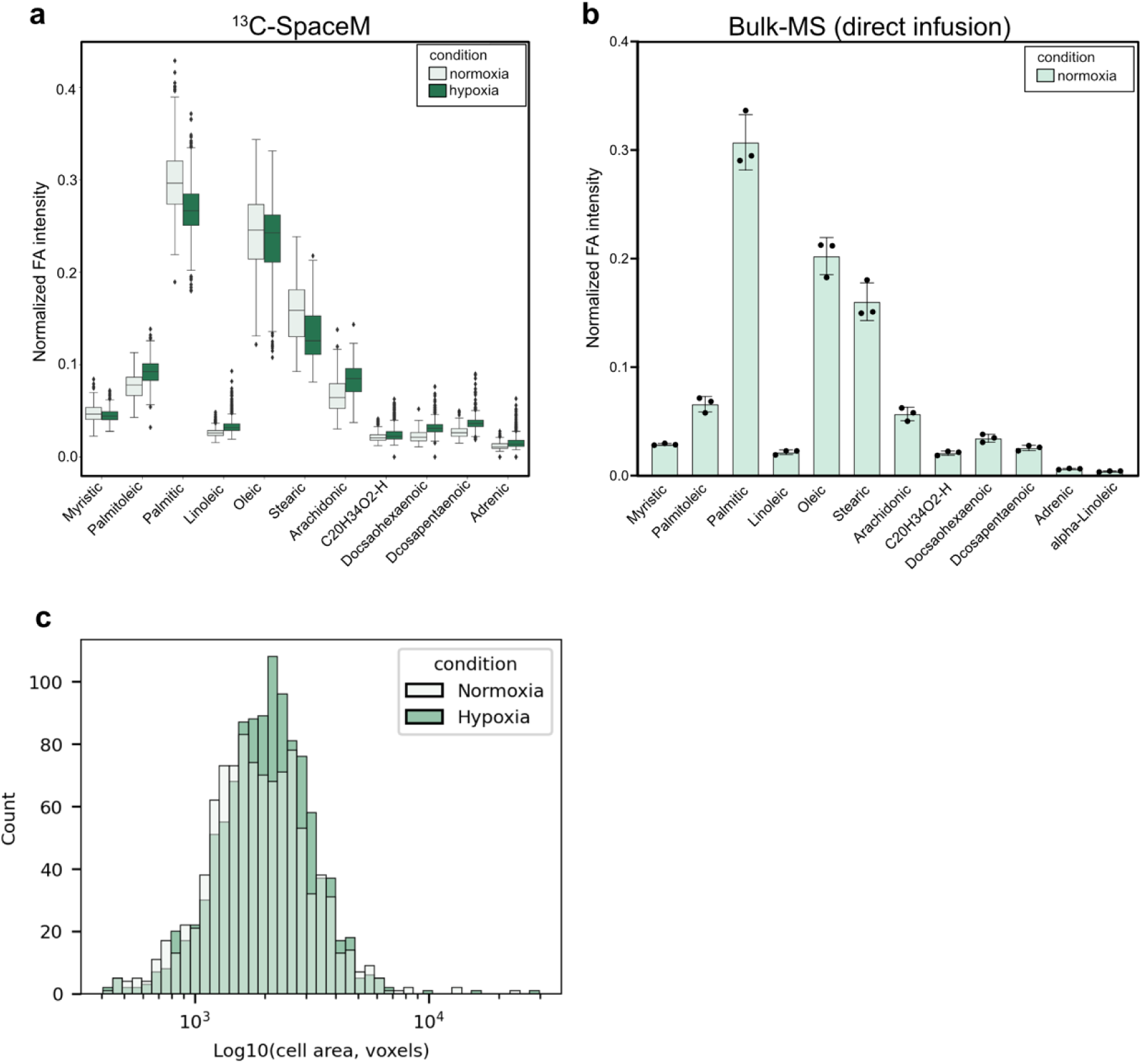
**a)** Box plots displaying mean and distribution of normalized intensities of 11 different fatty acids detected by 13C-SpaceM in normoxic and hypoxic cells. **b)** Normalized intensities of 12 fatty acids released from total lipids isolated by saponification from normoxic cells and quantified using bulk-MS with direct infusion. (n=3) **c)** Histogram of the distribution of cell area as determined by imaging for cells cultured in normoxia and hypoxia.

**Figure S2 (accompanying Figure 2).**
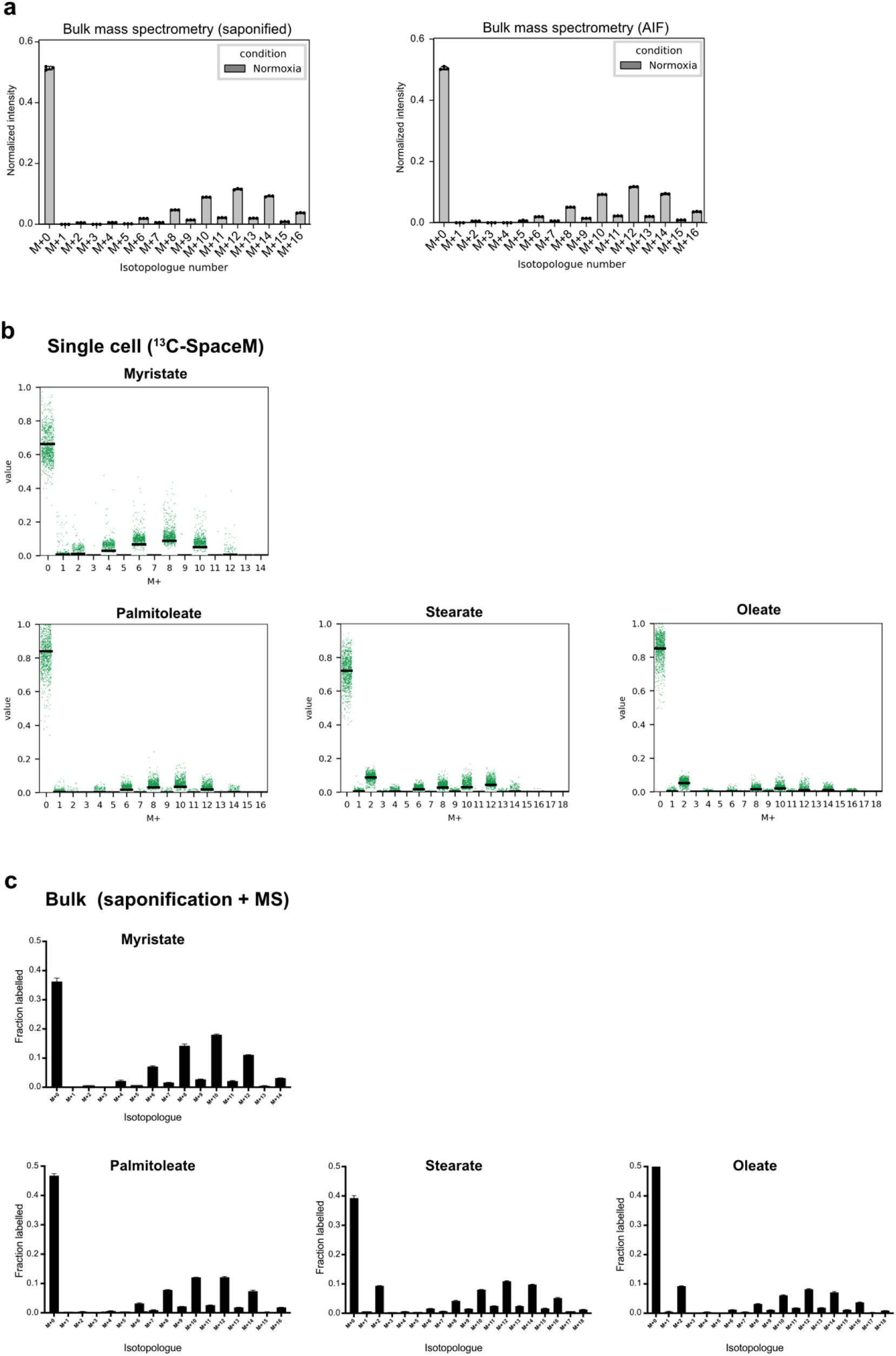
**a)** Comparison of isotopologue profiles of palmitate determined by bulk mass spectrometry after saponification (left graph) and all ion fragmentation (right graph) from cells cultured in normoxia. (n=3) **b)** Single cell isotopologue profiles for myristate, palmitoleate, stearate and oleate derived using ^13^C-SpaceM from cells cultured in normoxia. Black lines show average values. **c)** Isotopologue profiles for myristate, palmitoleate, stearate and oleate derived using bulk MS after saponification from cells cultured in normoxia. (n=3)

**Figure S3 (accompanying Figure 3):**
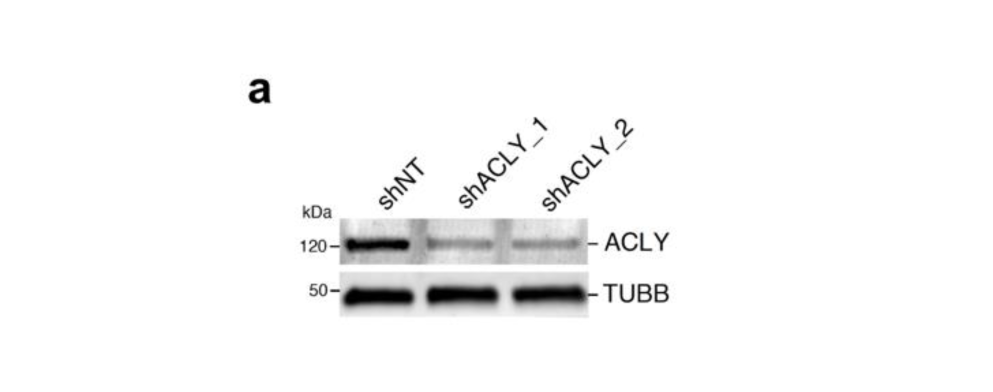
**a)** Immunoblot of protein lysates showing efficiency of ACLY knockdown using two shRNA sequences (ACLYkd oligo1 and ACLYkd oligo 2) compared to non-targeting control (shNT). Tubulin B (TUBB) is shown as loading control.

**Figure S4 (accompanying Figure 5):**
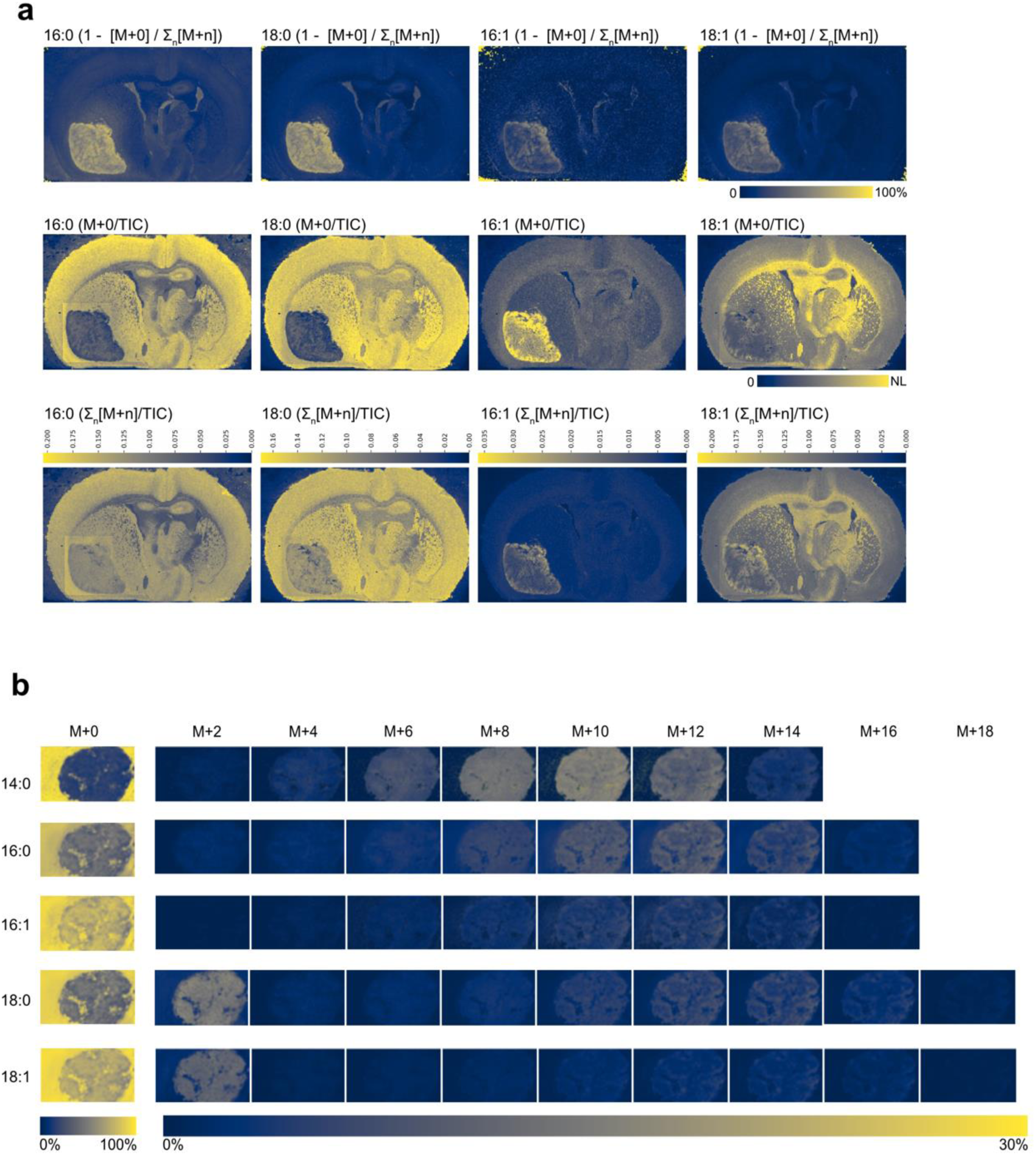
**a)** Top row: Fractions of palmitate, palmitoleate, stearate and oleate derived from fatty acid synthesis during the labelling period (48hrs) visualized by displaying the sum of all labelled isotopologues as a fraction of the sum of all isotopologues (1 - [M+0] / Σn[M+n]). Middle row: the unlabeled isotopologue (M+0) for palmitate, palmitoleate, stearate and oleate normalized to the TIC. Bottom row: Sum of all isotopologues shown as a fraction of the TIC, scale bar shows fraction of TIC. **b)** Isotopologue fractional images for myristate (14:0), palmitate (16:0), palmitoleate (16:1), stearate (18:0) and oleate (18:1) in tumor tissue (tumor #2) from a mouse orthotopically implanted with GL261 glioma cells and fed with a liquid diet containing U-^13^C_6_-Glucose for 48 hours. Colors represent fraction of each isotopologue of the sum of the all isotopologues for each fatty acid.

